# Sensitivity of the human temporal voice areas to nonhuman primate vocalizations

**DOI:** 10.1101/2025.09.19.677258

**Authors:** Leonardo Ceravolo, Coralie Debracque, Thibaud Gruber, Didier Grandjean

**Affiliations:** University of Geneva, Geneva, Switzerland

## Abstract

In recent years, research on voice processing in the human brain, particularly the study of temporal voice areas (TVA), was dedicated almost exclusively to conspecific vocalizations. To characterize commonalities and differences regarding primate vocalization representations in the human brain, the inclusion of closely related nonhuman primates, namely chimpanzees and bonobos, is needed. We hypothesized that neural commonalities would depend on both phylogenetic and acoustic proximities, with chimpanzees ranking closest to Homo. Presenting human participants (N=23) with the vocalizations of four primate species (rhesus macaques, chimpanzees, bonobos and humans) and regressing-out relevant acoustic parameters using three distinct analyses, we observed within-TVA, sample-specific, bilateral anterior superior temporal gyrus activity for chimpanzee vocalizations compared to: all other species; nonhuman primates; human vocalizations. Within-TVA activity was also observed for macaque vocalizations. Our results provide evidence for subregions of the TVA that respond principally, but not exclusively, to phylogenetically and acoustically close nonhuman primate vocalizations, namely those of chimpanzees.

## Introduction

The study of cerebral mechanisms underlying speech and voice processing has gained importance since the early 2000s with the advent of functional magnetic resonance imaging (fMRI) [1]. Voice-sensitive areas, commonly referred to as “temporal voice areas” (TVA) or simply “voice areas”, have been highlighted along the upper, superior portion of the temporal cortex [2]. Since then, great efforts have been made to better characterize these TVA, with particular attention to their spatial division into functional subregions [3–5]. A fairly large body of literature points to the critical role of the TVA in voice perception and processing in healthy participants [4, 6–8] as well as in lesioned patients [9]. Subregions of the TVA have also been directly linked to social perception [10], vocal emotion processing [11, 12], voice identity [13, 14], and gender perception [15]. The developmental axis of voice processing has also been studied in infants, demonstrating the existence of the TVA and voice sensitivity in the human brain as early as 4 [16] and 7 [17] months of age, while the ability to respond specifically to the voice of their parents has been observed in fetuses in utero [18]. With the ongoing development of brain imaging and analysis techniques [19], it is realistic to expect successful, albeit noninvasive, fMRI results on task-related voice perception in utero in the near future. Along the evolutionary axis, evidence for TVA or, more generally, conspecific vocalization-sensitive brain areas has emerged primarily in dogs [20], monkeys [21, 22] (*Macaca mulatta*) as well as recently in common marmosets [23, 24] (*Callithrix jacchus*), raising the question of whether such specialized brain areas are species-specific [25] and to what extent human and nonhuman primates share neural mechanisms that enable them to preferentially process conspecific vocalizations [26]. However, less attention has been paid to paradigms in which animal vocalizations are presented to humans—see Fecteau and colleagues’ study on species-specificity in the human brain [25]. Human processing of animal vocalizations has been studied with both monkey and cat material, but no *specific* cross-species activations have been observed within the TVA with respect to either species [27]. Other studies have focused more specifically on phylogenetic distance and have included nonhuman ape (chimpanzee, *Pan troglodytes*) and ‘Old World’ monkey (rhesus macaque, *Macaca mulatta*) vocalizations as stimuli. Such studies failed to identify species-specific brain activations—despite correctly discriminating chimpanzee affective vocalizations [28]—and observed ambivalent results for below [28] vs. above [29] chance discrimination of macaque affective vocalizations by human participants. A recent exception is a study in which functionally homologous anterior TVA activity was observed in both humans and macaques: this region was indeed specific to macaque calls in the macaque’s anterior TVA, and specific to human voices in the anterior TVA of humans, but no macaque-specific activity was observed in the human TVA [30]. Additionally, TVA responded similarly in both species to the presentation of nonverbal auditory stimuli, potentially due to some level of acoustic similarity [30]. This sparse literature motivated the present study, which aims to investigate cross-species TVA activations in humans when asked to categorize vocalizations from phylogenetically—and acoustically-close and -distant—species while undergoing fMRI scanning. While studies on the TVA typically involve passive listening, we wanted to additionally have the means to compare nonhuman primate species recognition/categorization in humans. The importance of acoustic differences between species and more specifically acoustic distance, particularly through fundamental frequency variations [31, 32] was indeed of great interest. Acoustic distance—calculated using Mahalanobis distance with 16 acoustic parameters extracted from the stimuli, see Table S1—was in fact a determining parameter in assessing affective cues recognition in nonhuman primate calls by human participants [33]. In this study, affiliative chimpanzee—but not bonobo—calls were acoustically the closest to positive human voice stimuli, suggesting a distinct evolution of bonobo calls [33]. Bonobo vocalizations are of particular interest because this species is thought to have undergone evolutionary changes in their communication, in part due to a neoteny process involving acoustic modifications, and although they are as phylogenetically close to humans as chimpanzees—with an estimated separation with the *Homo* lineage only 6-8 million years ago [34]. Previous research has shown that bonobos have a shorter larynx—a valid predictor of a species’ mean fundamental frequency [35]—compared to chimpanzees, resulting in a higher fundamental frequency in their calls [31]. Such a difference has been demonstrated in juvenile bonobo calls compared to chimpanzee and human baby calls [36], arguing for a greater acoustic distance between bonobo calls and human or chimpanzee vocalizations. For these reasons, we included vocalizations from both Pan species (chimpanzees, *Pan troglodytes*; bonobos, *Pan paniscus*), as well as a phylogenetically more distant species (*Cercopithecidae*: rhesus monkeys), with an estimated separation with the *Homo* lineage dating back to 25 million years ago. Indeed, any claim of human ‘uniqueness’ for TVA selectivity remains on hold and should be tested in light of these closely related species. Using the same stimuli, we previously investigated the frontal mechanisms involved in the categorization of nonhuman primate vocalizations independently of a selection of low-level acoustic parameters [37], but the possibility that acoustic differences would affect, at the auditory level, the ability of human participants to recognize nonhuman primate calls should be thoroughly examined, as we did in the present study. As suggested by research mentioned above, monkey vocalizations are overall less likely to be identified compared to ape vocalizations due to both phylogenetic and acoustic differences. Therefore, our mechanistic hypothesis of the difficulty for humans to recognize bonobo calls is that frequencies of the human tonotopic map in the auditory cortex—adapted and adjusted to the frequencies of the human voice during evolution—would not be tailored to process the frequencies generated by bonobo calls. It would also be the case for macaque calls, while frequencies of chimpanzee calls—being closer to the range of human voice fundamental frequency [31, 33]—would be better represented in the human auditory cortex and therefore more easily processed and better identified by humans.

According to the literature mentioned so far and to the mechanistic hypothesis underlying the processing of chimpanzee as opposed to bonobo and/or macaque calls by human participants, we therefore predicted: (i) Bioacoustics: more acoustic proximity between human and chimpanzee vocalizations, whereas more distance would separate those of bonobos and macaques from the human voice [31, 33, 38]; (ii) Phylogeny: a selectivity of the superior temporal lobe—within the TVA—for the processing of vocalizations from the Pan taxon (chimpanzee, bonobo) but not *Cercopithecidae* (rhesus monkey) vocalizations [25, 28, 34, 36, 39]—while considering acoustic features that characterize the stimuli the best through a discriminant analysis.

## Results

### Behavior: general aspects

Our hypotheses involve a control of phylogeny through the inclusion of specific primate species calls as well as the selection of specific acoustic features. We programmed a task in which the vocalizations of each species were presented randomly, and for which the participants (N=23) had to specify to which species each stimulus corresponded. We therefore included equal numbers of trials (N=72) with human, chimpanzee, bonobo and macaque vocalizations—N=18 each—as well as trial-level acoustic features of the vocalizations, using distinct statistical models with specific covariates. The stimuli were presented at a constant sound pressure level but the stimuli were not normalized to avoid altering their naturality [40]. The statistical models are sorted from the least to the most sophisticated modeling to uncover the role(s) of acoustic features on TVA activity potentially specific to each/some species (see the Methods; Model 1: mean of vocalization fundamental frequency and energy; Model 2: multi-dimensional Mahalanobis acoustic distance between the human voice and the calls of each nonhuman primate species [41]; Model 3: between-species most discriminant acoustic features of our stimuli, extracted using a general discriminant analysis [33]; Model 4: model-based approach of the probability of correct species categorization, including the most discriminant acoustic features of Model 3). Acoustical analyses involved in Model 2 allowed us to validate our first hypothesis according to which chimpanzees are acoustically the closest to humans, followed by the calls of bonobos and macaques (Fig.1**C**)— the main effect of Species on the acoustic distance was significant, F(3,88)=15.84, *p*<.001, as well as all comparisons (Hum > Chimp: t(23)=-12.22, *p*<.001; Hum > Bon: t(23)=-14.16, *p*<.001; Hum > Mac: t(23)=-22.57, *p*<.001; Hum > (Chimp,Bon,Mac): t(23)=-20.86, *p*<.001; Chimp > Bon: t(23)=-5.07, *p*<.01; Bon > Mac: t(23)=-6.43, *p*<.001; Chimp > (Bon,Mac): t(23)=-27.34, *p*<.001; see also Table S2).

**Fig. 1:**
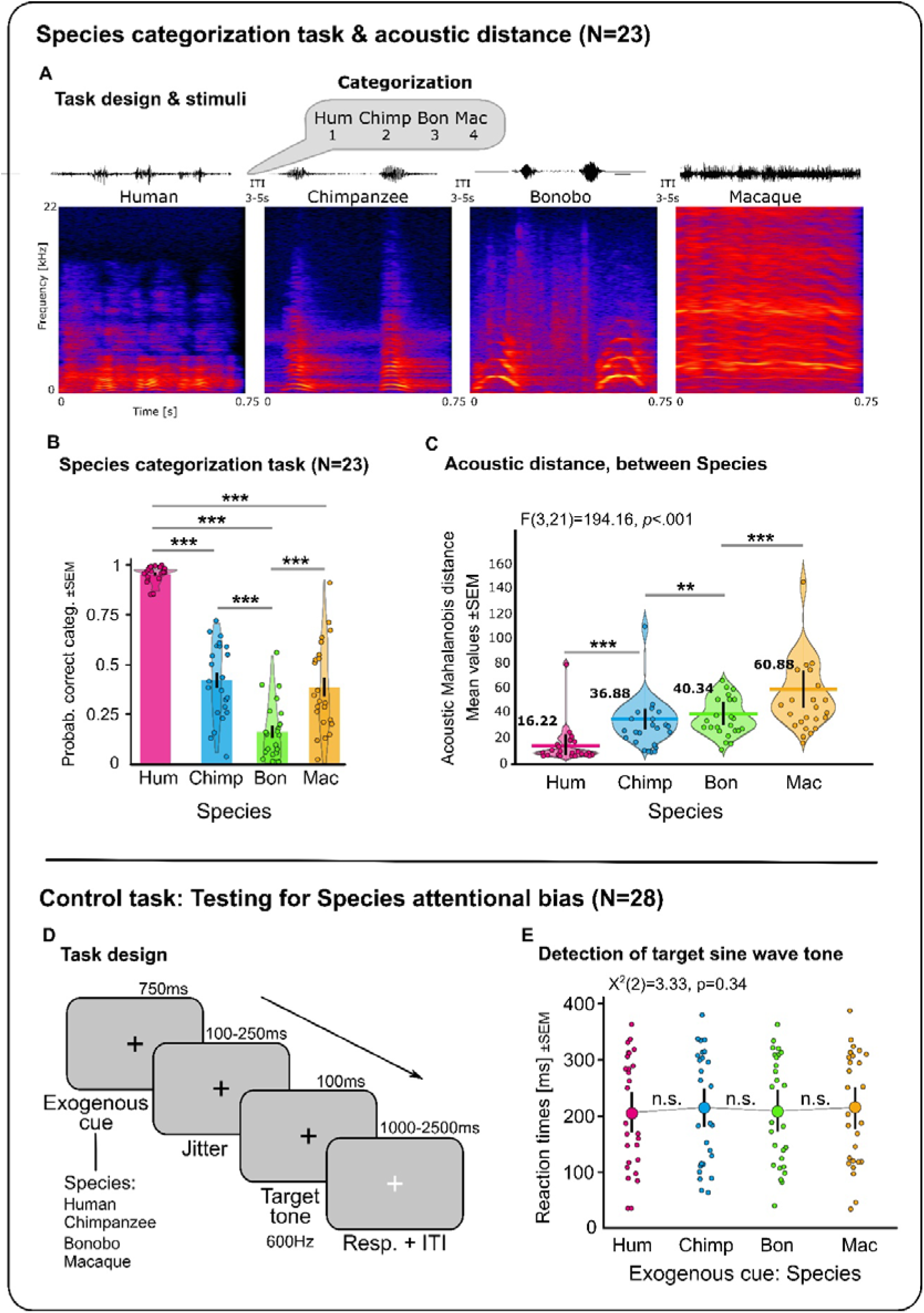
Timecourse and results of the species categorization task and the control task testing for potential attentional biases. (**A**) Detail of the timecourse of four trials of the species categorization task in non-representative order, including waveform and spectrogram graphs for one example stimulus of each species. (**B**) Behavioral results of the task (N=23) showing the probability (Probab.) of correctly categorizing (categ.) each Species’ vocalization, using the 6 acoustic covariates from Model 3 and reaction times as variables. (**C**) Histograms of the acoustic Mahalanobis distance data of each species including mean (numbers represent exact mean value). (**D**) Control task and its design in an independent sample of 28 participants using each species’ vocalization as an exogenous cue preceding the target to be detected as fast as possible, namely a short sine wave tone (600Hz). (**E**) Results of the control task (N=28), showing no biasing of attention by any of the species stimuli: no species triggered attentional capture, yielding to no attentional advantage for target detection. For the results plots, violin plots illustrate distribution fit, error bars the standard error of the mean (SEM). Points represent individual values. ITI: inter trial interval; Resp.: response; Hum: human; Chimp: chimpanzee; Bon: bonobo; Mac: macaque. \*\**p*<.01, \*\*\**p*<.001, n.s.: non-significant.

### Behavioral data: species categorization task

As described above, Model 4 was computed to analyze the ability of our participants to categorize the species vocalizations, considering their production social contexts, namely agonistic (threat, distress) or affiliative (‘positive’). These analyses used as additional predictors the six most discriminant acoustic features described in Model 3, in addition to controlling for response reaction times (Fig.1**A**,**B**). These data revealed a main effect of the Species and Context factors (χ^2^(3)=187.35, *p*<.0001 and χ^2^(2)=6.62, *p*<.05) as well as their interaction (χ^2^(6)=14.02, *p*<.05). Main effects of acoustic features were also significant: F0 power (χ^2^(1)=4.44, *p*<.05) and intensity contour difference (χ^2^(1)=6.07, *p*<.05). Species categorization results were explained by human voices being categorized more accurately than all other species (human vs nonhuman primate species: χ^2^(1)=144.31, *p*<.0001; human vs chimpanzee: χ^2^(1)=97.09, *p*<.0001; human vs bonobo: χ^2^(1)=177.91, *p*<.0001; human vs macaque: χ^2^(1)=98.34, *p*<.0001) and by worse, below chance-level categorization of bonobo calls compared to all other species (bonobo vs human: χ^2^(1)=177.91, *p*<.0001; bonobo vs chimpanzee: χ^2^(1)=22.86, *p*<.0001; bonobo vs macaque: χ^2^(1)=28.84, *p*<.0001). Interestingly, no difference in accuracy was observed for the categorization of chimpanzee compared to macaque calls (χ^2^(1)=0.004, *p*=.94). Since Context was controlled for but not part of our hypotheses, and since it was strongly interacting with Species, we report all posthoc contrasts in Table 1.

**Table 1:**
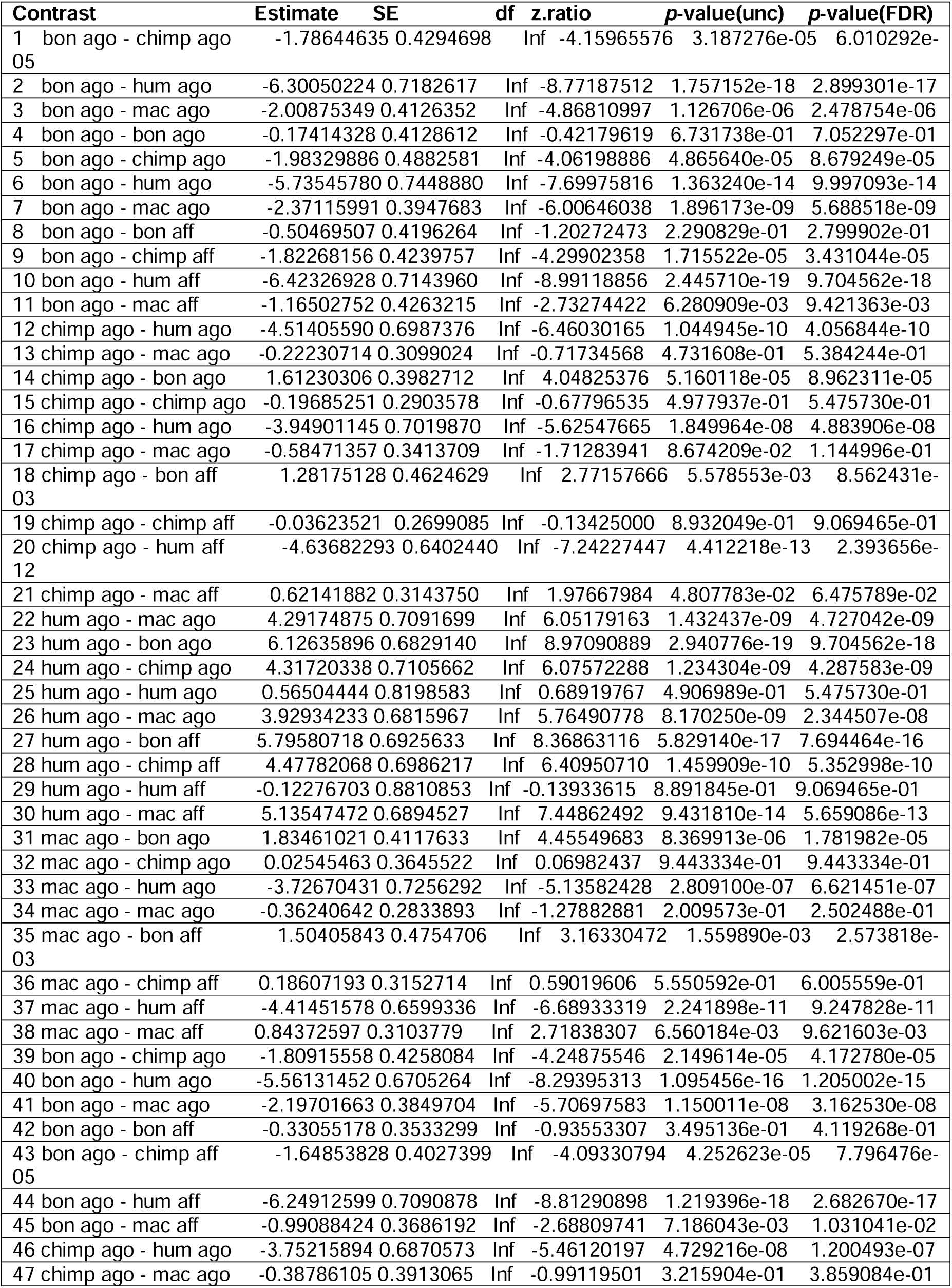
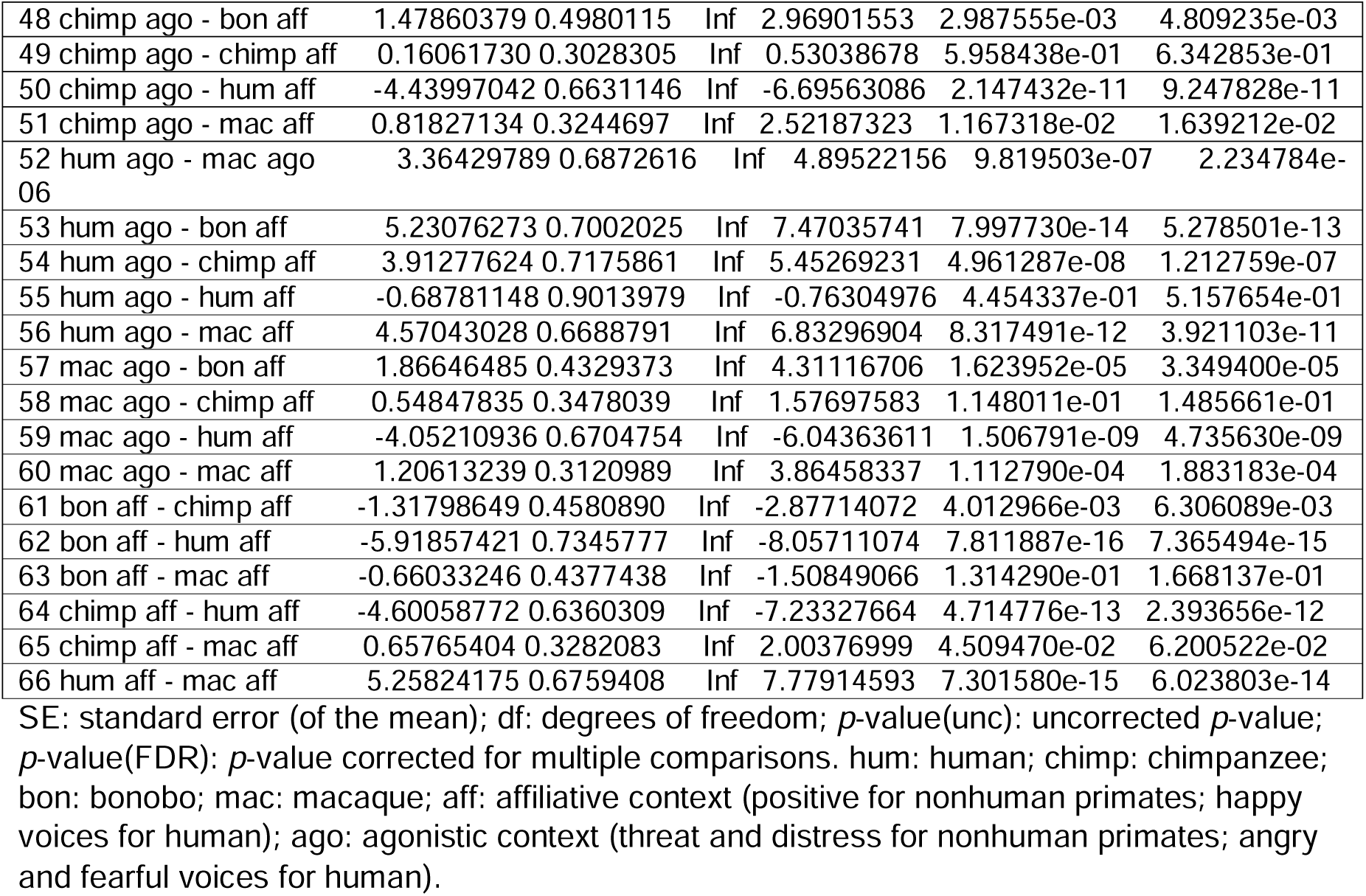
Posthoc contrasts for behavioral effects of the Species by Context interaction.

**Table 2:** Activations, cluster size and coordinates for each contrast of interest of model 3 (vocalization loudness, intensity, chagoe in spectrum, F2 bandwidth contour, F0 power and intensity contour difference as trial-level covariate of no-interest) in the sample-specific temporal voice areas, wholebrain voxelwise *p*<.05 FDR corrected, k>10.

| MNI coordinates |  |  |  |  |  |  |
| --- | --- | --- | --- | --- | --- | --- |
| Region label<br>(voxels) | Hemisphere | X | Y | Z | T value | Cluster size |
| **Chimpanzee > Human, Bonobo, Macaque** |  |  |  |  |  |  |
| Superior temporal gyrus ant<sup>10</sup> | L | -52 | -2 | -12 | 4.54 | 72 |
| Superior temporal gyrus ant<sup>11</sup> | R | 54 | 0 | -12 | 3.19 | 19 |
| **Chimpanzee > Bonobo, Macaque** |  |  |  |  |  |  |
| Superior temporal gyrus ant<sup>13</sup> | L | -54 | -2 | -12 | 5.03 | 100 |
| *Superior temporal gyrus ant* | L | -58 | -12 | -2 | 4.98 |  |
| *Superior temporal gyrus ant* | L | -52 | 2 | -12 | 4.38 |  |
| Superior temporal gyrus ant<sup>14</sup> | R | 58 | -2 | -12 | 4.97 | 91 |
| *Superior temporal sulcus ant* | R | 54 | -8 | -12 | 4.30 |  |
| **Chimpanzee > Human** |  |  |  |  |  |  |
| Superior temporal gyrus ant<sup>12</sup> | L | -50 | -4 | -12 | 2.70 | 10 |
| *Superior temporal gyrus ant* | L | -48 | -10 | -12 | 2.60 |  |
| **Human > Chimpanzee** |  |  |  |  |  |  |
| Superior temporal gyrus mid | L | -54 | -14 | 4 | 8.55 | 5392 |
| *Supramarginal gyrus* | L | -60 | -48 | 24 | 7.54 |  |
| *Superior temporal gyrus post* | L | -54 | -58 | 18 | 6.88 |  |
| *Middle temporal gyrus mid* | L | -66 | -22 | -8 | 6.39 |  |
| *Middle temporal gyrus post* | L | -66 | -44 | -2 | 5.49 |  |
| Supramarginal gyrus | R | 56 | -42 | 28 | 8.22 | 5250 |
| *Supramarginal gyrus* | R | 56 | -42 | 36 | 7.52 |  |
| *Superior temporal gyrus mid* | R | 56 | -8 | 2 | 7.50 |  |
| *Superior temporal gyrus post* | R | 50 | -48 | 22 | 6.23 |  |
| *Superior temporal gyrus post* | R | 58 | -46 | 14 | 6.22 |  |
| *Superior temporal gyrus mid* | R | 50 | -18 | 6 | 6.05 |  |
| *Superior temporal gyrus post* | R | 60 | -46 | 18 | 5.99 |  |
| *Middle temporal gyrus post* | R | 70 | -32 | 0 | 5.84 |  |
| *Middle temporal gyrus mid* | R | 64 | -22 | -12 | 5.75 |  |
| *Middle temporal gyrus mid* | R | 66 | -20 | -18 | 5.42 |  |

ant: anterior; mid: central part; post: posterior.

These results cannot be attributed to potential attentional biases towards or saliency between the calls/vocalizations of our species. Indeed, no difference in exogenous cueing was observed in an independent sample of 28 participants in a dedicated task (see Methods and Fig.1**D**,**E**).

### Neuroimaging data within the sample-specific temporal voice areas

We aimed at uncovering functional changes relative to species categorization and processing within sample-specific (N=23) TVA, as delineated in our hypotheses. As described above, we used three distinct and independent statistical models including trial-level parametric modulators (Model 1-3). We were particularly interested in human brain activity while processing vocalizations of our closest relatives—both acoustically and phylogenetically—namely the chimpanzee but also the bonobo. The present study did not aim at uncovering wholebrain results underlying the processing of each species’ vocalizations but rather focused on human voice-sensitive areas, namely the TVA, although corrected statistics (voxelwise *p*<.05 False Discovery Rate) presented in this section were computed with a wholebrain voxelwise approach for higher data reproducibility and generalizability, and not using region-of-interest (ROI) analyses. ROI analyses would most probably have artificially amplified the number of voxels in the TVA in this study. Clusters outside the bounds of the sample-specific TVA are therefore visible but in a desaturated hue to better highlight TVA activations. These clusters are more visible in supplementary figures with the same contrasts as those presented in this section, but with an outline of the TVA from an independent, larger sample of participants excluding the 23 participants of this study (N=98; Fig.S1-3).

The three statistical models tested the impact of literature-based [31, 33, 34, 36] acoustics (1), acoustic distance between our species (2) and data-driven acoustics that represent our stimulus set the best (3). Results from all models overlap greatly, showing within-TVA enhanced activity mostly for chimpanzee calls compared to the other primate vocalizations—both when including or excluding human voice stimuli. In this section, we will focus on the latter model (3), which is the most sensitive and therefore the most meaningful for the present study. Activations coordinates can be found in Table 1, illustration in Fig.2. Data from Model 1-2 can be found in the supplementary material, see Fig.S1-3 and Table S3-4.

**Fig. 2:**
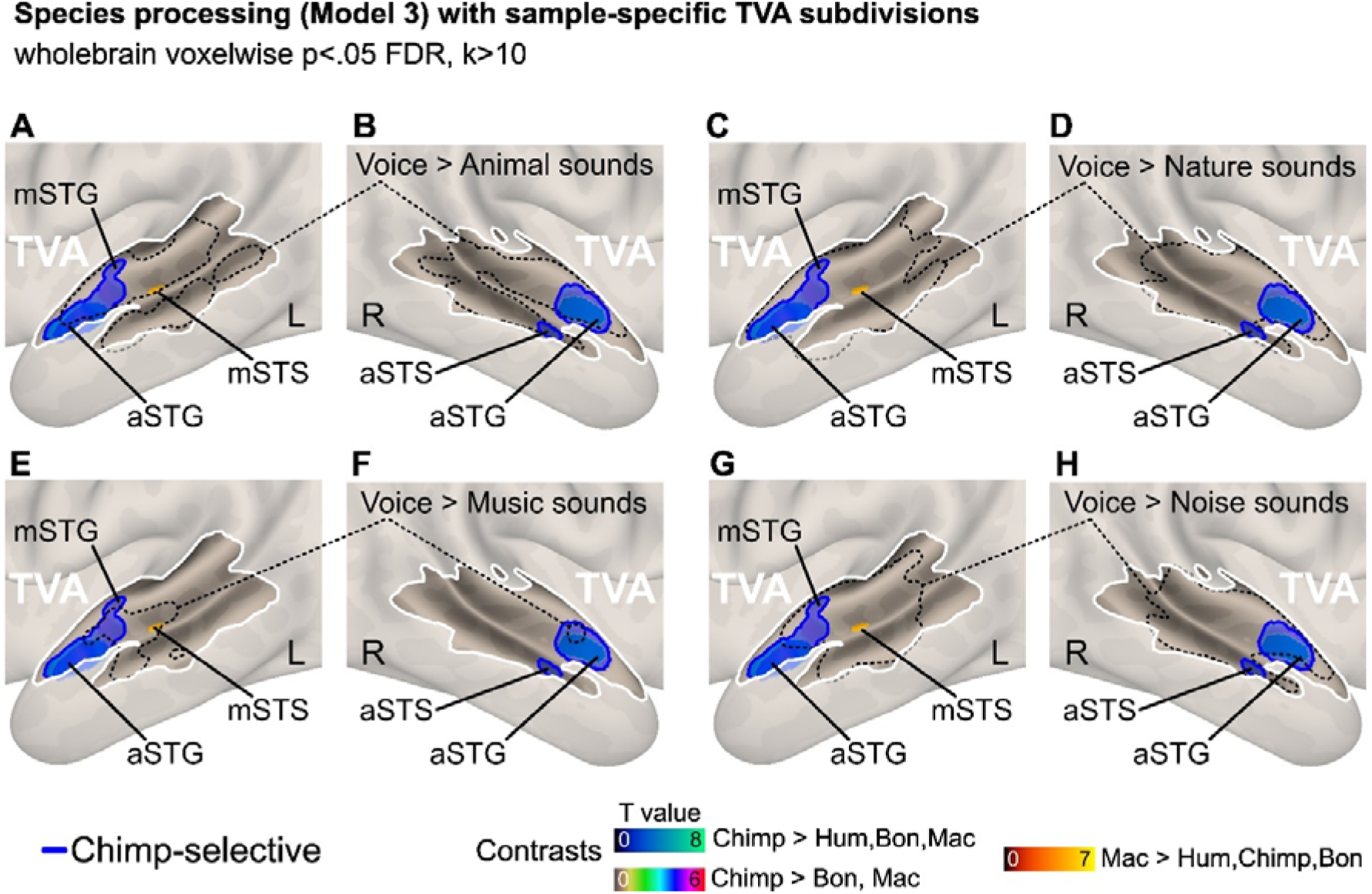
Wholebrain results with TVA outlines when contrasting the processing of chimpanzee to other species’ vocalizations with vocalization loudness, intensity, change in spectrum, F2 bandwidth contour, F0 power and intensity contour difference as trial-level covariates of no-interest (model 3). Enhanced brain activity on a sagittal view with activity for [chimpanzee > human,bonobo,macaque] (dark blue to green), [chimpanzee > bonobo,macaque] (brown to red with light yellow outline) and [macaque > human,bonobo,chimpanzee] (red to yellow) vocalizations, with outlines of TVA for voice > animal sounds (**A,B**), voice > nature sounds (**C,D**), voice > music (**E,F**) and voice > noise (**G,H**). Brain activations are independent of the most discriminant low-level acoustic parameters of the stimuli set [33]. Data corrected for multiple comparisons using wholebrain voxelwise false discovery rate (FDR) at a threshold of *p*<.05. Hum: human; Chimp: chimpanzee; Bon: bonobo; Mac: macaque. White outline: sample-specific temporal voice areas (TVA; N=23); Dotted black outline: sample-specific TVA, per sound category; Blue outline: areas selective to chimpanzee calls. ‘a’ prefix: anterior; ‘m’ prefix: mid; ‘p’ prefix: posterior; STG: superior temporal gyrus; STS: superior temporal sulcus; L: left hemisphere; R: right hemisphere.

In addition to the abovementioned models, we also included a model-based methodology (Model 4) to uncover the brain correlates of the probability of correctly categorizing the four species, within the sample-specific TVA. This modeling includes the fitted regression coefficients computed using as covariates the acoustic parameters of Model 3, in addition to task reaction times. These neuroimaging results are presented in Fig.4.

### Effects of species processing with vocalization most discriminant acoustic parameters (N=6) as covariates of no-interest at the trial level

In this third model, we wanted to elaborate more on the discriminant factors that characterize the low-level acoustic parameters of our set of stimuli. This approach is complementary to the inclusion of both basic acoustics (model 1) and acoustic distance in model 2 and therefore refines these results. To do so, we used as trial-level covariates of no-interest the acoustic parameters explaining the most variance ([r > 0.7] and [r < -0.7]) in factors 1-3 of a discriminant analysis of these stimuli [33]—see the methods section for details on this analysis. These parameters therefore include, in this specific order: vocalization loudness, intensity, change in spectrum, bandwidth contour of the second formant (F2), power of the fundamental frequency (F0) and finally the difference in intensity contour. We used these acoustic features as covariates in our data modeling. As in previous modeling of the imaging data, TVA activity was triggered by chimpanzee vocalizations ([chimpanzee > human, bonobo, macaque]) in large bilateral clusters of the aSTG within the TVA (Fig.2**A****-H**), closely resembling activations of model 2 (Fig.S2). Chimpanzee compared to other nonhuman primate calls in this model ([chimpanzee > bonobo, macaque]) led to the largest clusters observed in the aSTG—all models considered, still within the sample-specific TVA. Indeed, we observed a large left-lateralized cluster of the aSTG extending to the mid STG (Fig.2**A****,C,E,G**) as well as a right-lateralized cluster (Fig.2**B****,D,F,H**). These chimpanzee-selective areas are outlined in blue in each panel of Fig.2.

In this last modelling of the fMRI data, no voxels reached significance either at the wholebrain level or within the TVA for the [bonobo > human, chimpanzee, macaque] or [bonobo > chimpanzee, macaque], [bonobo > chimpanzee], [bonobo > macaque] contrasts. We however found activity for the processing of macaque calls only in the left TVA, more specifically in a small cluster of the left mid STS ([macaque > human, chimpanzee, bonobo], Fig.2**A****,C,E,G**)—and in a small portion of the planum temporale adjacent to the primary auditory cortex for the [macaque > chimpanzee, bonobo] contrast (see Fig.S4). Other contrasts are reported in Fig.S5.

### A synthesis of the sensitivity of the human TVA to nonhuman primate calls (across statistical models)

In the previous sections, we described three different models used to analyze our fMRI data. These models—from the simplest to the more sophisticated one—highlighted enhanced activity within sample-specific bilateral anterior TVA of our participants specifically when processing chimpanzee vocalizations—but also when processing macaque calls in model 3, in the bilateral mid STG, STS and planum temporale. When processing chimpanzee calls, TVA activity was especially enhanced in the aSTG but also in the anterior STS. We therefore regrouped these fourteen chimpanzee-selective aSTG clusters in Fig.3—most of them overlap greatly but we still named them individually according to each contrast and analysis for exhaustivity—overlaid with sample-specific TVA (Fig.3**C****,D**) and with the more general TVA from an independent sample of ninety-eight participants (Fig.3**A****,B**). Zooming closely, the area of maximal overlap between these regions (the orange surface) is located within the more general as well as within the sample-specific TVA. Interestingly, left-lateralized more medial clusters of aSTG were outside the outline of the sample-specific but not of the general TVA (Fig.3**A****,C**), while this was not the case for right-lateralized aSTG activations. Comparing the areas recruited when processing chimpanzee to bonobo and macaques calls, this contrast—especially in model 3, yielded to distinct clusters of aSTG. This result is visible when looking at the three ‘rich blue’ outlines in every panel of Fig.3. The results synthesized here highlight the important role of acoustic parameters and emphasize the role of the most discriminant acoustic features on TVA activity relating to nonhuman primate vocalizations, especially those of chimpanzees and macaques.

**Fig. 3:**
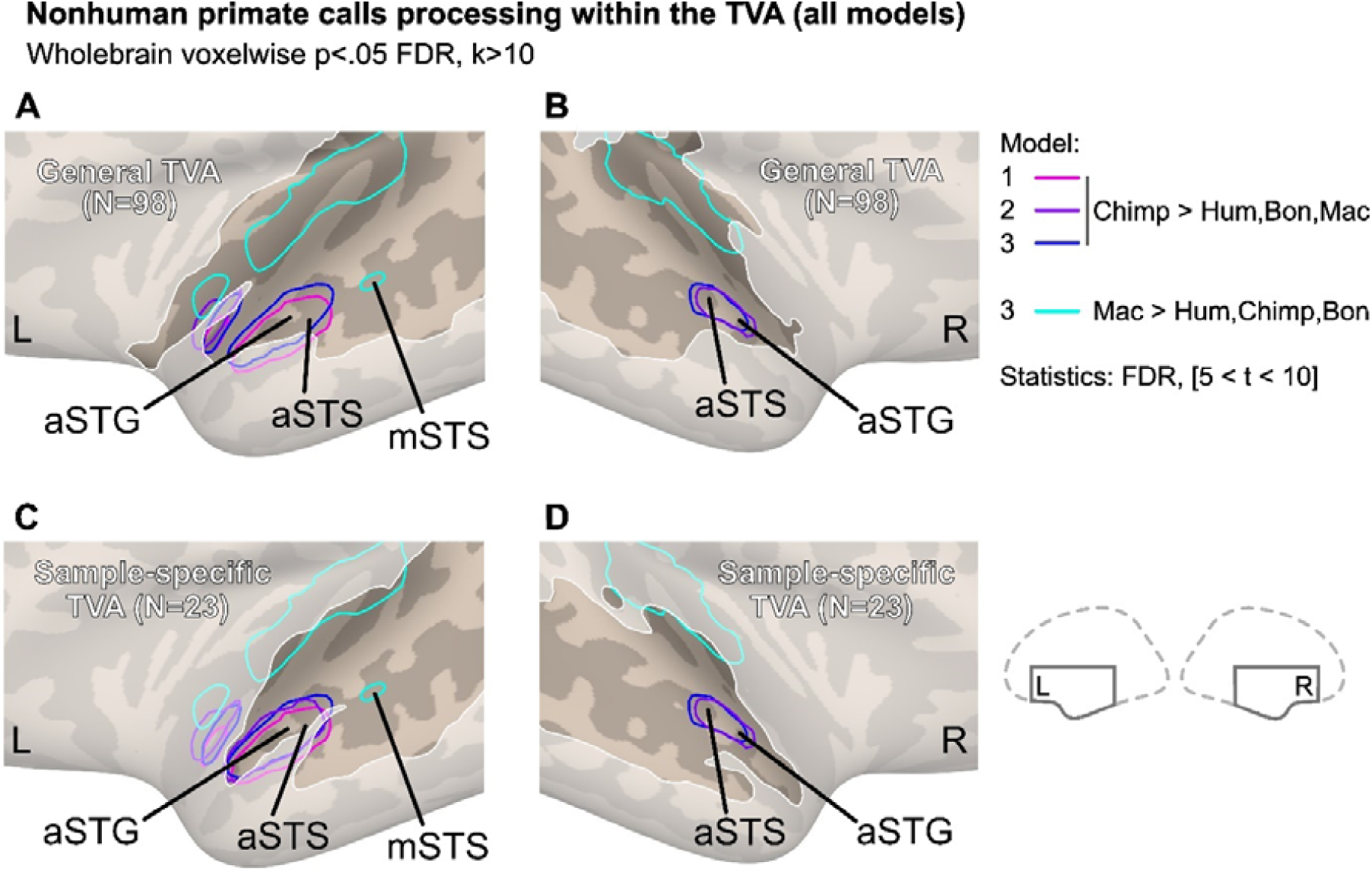
Synthesis of mid-to-anterior TVA clusters of activity recruited specifically by the processing of chimpanzee and macaque vocalizations (Models 1,2,3). Anterior superior temporal gyrus (aSTG) and sulcus (aSTS) clusters recruited for the processing of chimpanzee calls as opposed to human voices, bonobo, macaque calls (pink: model 1; purple: model 2; blue: model 3) in the general TVA (**A,B**, N=98) as well as in the sample-specific TVA (**C,D**, N=23). Macaque results are only significant for Model 3 (teal: Macaque vs all other species). Model 1: mean of fundamental frequency and energy (covariates of no-interest, N=2); Model 2: acoustic distance (covariate of no-interest, N=1); Model 3: acoustic parameters that characterize low-level acoustics of our stimuli following a discriminant analysis (covariates of no-interest, N=6). Data are all corrected for multiple comparison using wholebrain voxelwise false discovery rate (FDR) at a threshold of *p*<.05 with t-values ranging from 5 to 10. Hum: human; Chimp: chimpanzee; Bon: bonobo; Mac: macaque. TVA: temporal voice areas. Prefix ‘a’: anterior; ‘m’: mid. L / R: left / right hemisphere.

These results are given even more weight by more fined-tuned comparisons of voice versus non-voice material in the voice-localizer task, namely by splitting the non-vocal blocks as a function of the auditory sounds they contain. In this more specific outline of TVA subregions, visible in Fig.2 (as defined in Fig.S4) we observed that most chimpanzee- and macaque-selective STG and STS regions were still within the bounds of TVA.

### Within-TVA neural correlates of the probability of correct species categorization

Determining the neural correlates of the probability of correct species categorization within the sample-specific TVA was done using a model-based approach. Specific brain correlates of behavioral data on species categorization in frontal brain regions—using different acoustic covariates modeling—can be found elsewhere [37].

The present data revealed massive TVA activations when including all species together (including Human), namely in the posterior, mid and anterior STG, STS and middle temporal gyrus (MTG; Fig.4**A****,B**). Looking separately at these neural correlates for each nonhuman primate species, results show smaller, more specific areas for chimpanzee vocalizations, namely bilateral mSTG and mSTS in addition to right mMTG (Fig.4**C****,D**). For bonobo vocalizations, large correlates were observed bilaterally, covering most of the TVA. More specifically, the strongest peaks were located in the bilateral mSTG, mSTS and mid and posterior MTG (Fig.4**E****,F**). Finally, for macaque vocalizations, correlates were solely observed in a small cluster of the right mSTS (Fig.4**G****,H**). According to these data, correctly categorizing our species stimuli was associated with strong and vast TVA regions for bonobo calls, corresponding to the species with the lowest accuracy of all—and the only species with below-chance average values.

**Fig. 4:**
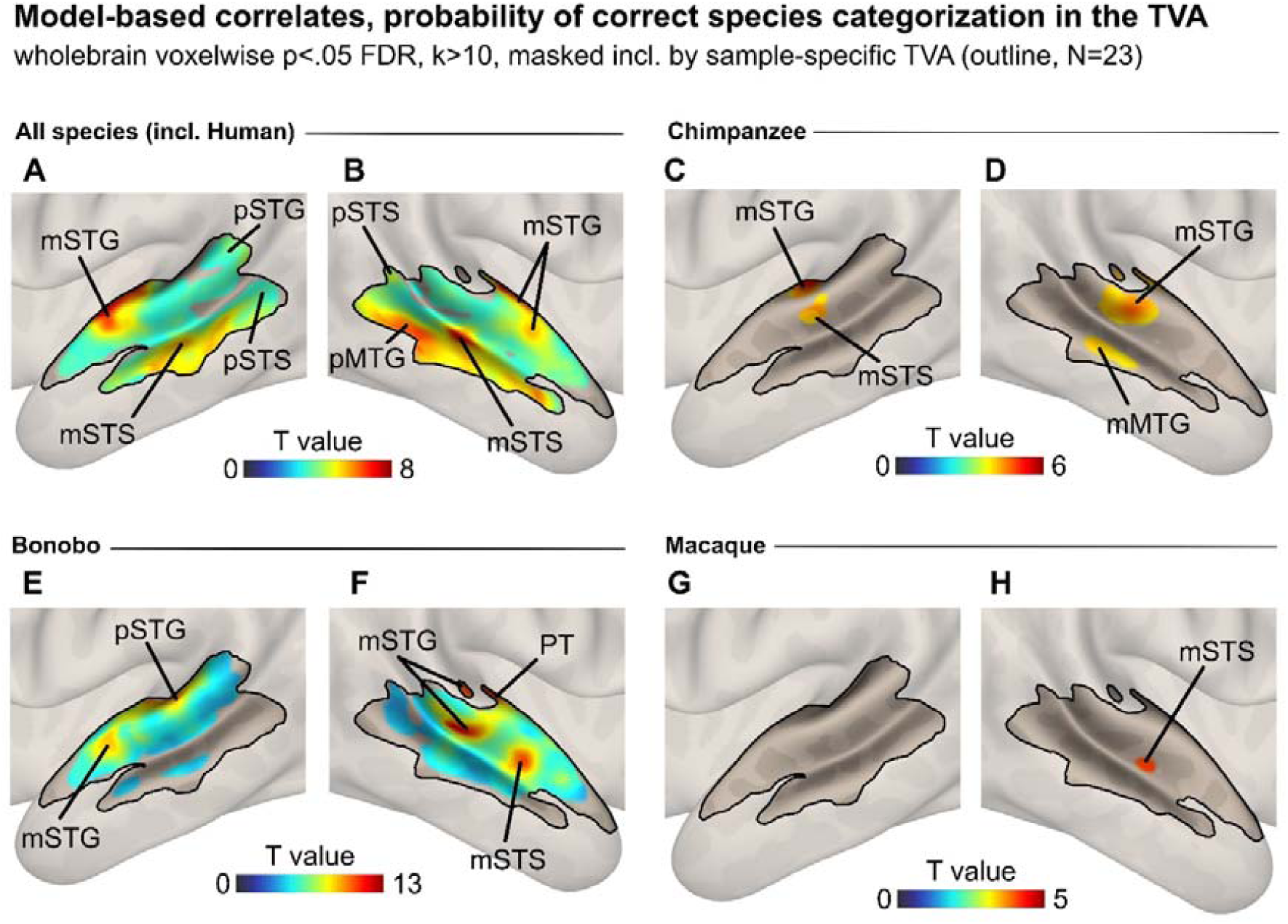
Model-based correlates of the probability of correct species categorization, within sample-specific TVA (Model 4). Correlates of the probability of correct species categorization computed using model-based analysis technique for all species, as illustrated on sagittal renders for all species, including human (**A,B**), then specifically for chimpanzee calls (**C,D**), bonobo calls (**E,F**) and macaque calls (**G,H**). These correlates were constrained to the bounds of the sample-specific TVA (N=23, black outline) using an inclusive masking procedure with correction for multiple comparison using voxelwise false discovery rate (FDR) at a threshold of *p*<.05. The colorbars represents T-value statistics. TVA: temporal voice areas. Prefix ‘a’: anterior; ‘m’: mid; ‘p’: posterior. STG: superior temporal gyrus; STS: superior temporal sulcus; MTG: middle temporal gyrus; PT: planum temporale. L / R: left / right hemisphere.

### Acoustic differences between chimpanzee and bonobo calls

The differences observed between chimpanzee and bonobo stimuli, both at the behavior and brain levels, may originate in their acoustic specificities—while phylogeny relative to humans is almost identical. While we will discuss this aspect in the Discussion, we explored it using a separate acoustic analysis dedicated to highlighting such acoustic differences. This analysis is different from the one in Model 3—although the same methods were used, because here we only use as input for GeMAPS [42, 43] the calls from chimpanzees and bonobos, therefore constraining acoustic characterization to these two species. The results highlight the acoustic parameters that explain the most variance for these specific species (see Methods and Table S5). The reduced parameters explaining at least 80% of the acoustic variance between chimpanzee and bonobo calls relate to (in that order): (1) pitch variation within the call (‘F0semitoneFrom27.5Hz_sma3nz_pctlrange0.2’), (2) within-call variability in mid-frequency range energy, sensitive to changes in voice quality and resonance (‘slopeV500.1500_sma3nz_stddevNorm’), (3) vocal tract opening (‘F1frequency_sma3nz_stddevNorm’), (4) irregularity in the temporal structure of unvoiced intervals (‘StddevUnvoicedSegmentLength’), (5) ratio of periodic to aperiodic energy (voice regularity, breathiness; ‘HNRdBACF_sma3nz_stddevNorm’), (6) spectral slope between 0 and 500 Hz in voiced frames (‘creaky’ or ‘pressed’ phonation; ‘slopeV0.500_sma3nz_amean’), (7) instability in glottal closure quality (‘F1bandwidth_sma3nz_stddevNorm’), (8) steepness of pitch rise between consecutive voiced frames (‘F0semitoneFrom27.5Hz_sma3nz_stddevRisingSlope’), (9) changes in place of articulation or lip posture (‘F2amplitudeLogRelF0_sma3nz_stddevNorm’), (10) within-call changes in spectral shape of very low frequency (‘slopeV0.500_sma3nz_stddevNorm’), (11) lip rounding and vocal tract length (‘F3amplitudeLogRelF0_sma3nz_stddevNorm’), (12) Jitter variability (micro-irregularities in vocal fold vibration; ‘jitterLocal_sma3nz_stddevNorm’), (13) spectral tilt measure (within-call changes in voice quality; ‘logRelF0.H1.A3_sma3nz_stddevNorm’).

## Discussion

The present study provides evidence of the sensitivity of the human TVA to cross-species vocalizations, especially to chimpanzee calls but also to macaque vocalizations, as illustrated by selective enhanced activity in the bilateral mid and anterior STG and STS—within sample-specific TVA. We do not interpret these results as ‘absolute’ selectivity, but rather we emphasize the relative preference of these brain regions to chimpanzee and to a lesser extent macaque calls. These results were obtained through statistical modeling of the MRI data that included either simple acoustics or the use of Mahalanobis acoustic distance between species and the most discriminant acoustic features specific to our stimuli as covariates. These two latter analyses converged and yielded to greatly overlapping results, especially in the anterior TVA. Therefore, our results suggest that vocalizations from another ape species recruits, or triggers activity in, subregions of human temporal cortex that process species-specific voices in humans—namely the bilateral, sample-specific TVA. This evidence speaks in favor of cross-species primate vocalization processing in the anterior and mid TVA of humans—for chimpanzee and macaque calls, respectively. Our results also suggest differential engagement of the TVA according to behavioral performance for species categorization. While our acoustic data confirmed the hypothesized hierarchy of acoustic distance as a function of phylogenetic distance between our species, we still observed mid STG and STS activity for macaque versus bonobo calls and a small cluster in the left mid STS selective to macaque calls in model 3—an unexpected result since macaques are the most distant species from humans both phylogenetically and acoustically in our study. Therefore, while we initially hypothesized that primate calls would exclusively enhance activity in human TVA as a function of a combination of phylogenetic and acoustic proximity, our data also point toward the greater importance of the most discriminant acoustic features rather than acoustic distance alone. We discuss these aspects below in more detail and interpret their general meaning and subsequent scientific implications, in addition to highlighting the limitations of our study.

Often specifically associated with the processing of conspecific vocalizations (e.g., in humans [2, 25, 30], macaques [22, 30, 44], and dogs [20]), the present study challenges the common view of the TVA as ‘species-specific’ and illustrates that human voices, chimpanzee and macaque calls can enhance activity in the TVA. We think that the distinct locations of these TVA subregions recruited for processing the vocalizations of these primate species matters. In fact, there might be a possible association between anterior TVA—selective to processing chimpanzee calls—and the higher recognition performance of chimpanzee calls compared to those of bonobos or macaques in human participants [37]. Anterior TVA activity selective to the processing of chimpanzee calls occurred when these were compared to both human and nonhuman primate species, solely to other nonhuman primate vocalizations, or directly to the human voice. However, homologous results were not observed for bonobo and they were more scarce—especially between models—for macaque vocalizations: we found macaque-selective activity in a small area of the planum temporale and in a small cluster of the left mid STS, congruent for instance with locations observed in the general processing of animal sounds, especially in the planum temporale [45, 46]. On the other hand, within-TVA anterior STG activity was also observed when chimpanzee vocalizations were directly compared to the human voice. We think this result highlights the cross-species selectivity of this anterior subregion of the TVA for processing species phylogenetically close to humans and especially with human-like acoustics, namely the calls of chimpanzees in the case of our study. Because of their vocal proximity, the perception of human voices and chimpanzee calls in socio-affective contexts could involve a common ‘social’ core of the brain, which increases activity in brain regions such as the anterior TVA, as reported previously in studies pertaining to social contextual information processing in the anterior STG [47, 48]. Differences at the level of processing complexity between the two types of vocalizations could also explain this observation, while we demonstrated that saliency or attention-related effects do not exist between our species stimuli. Indeed, previous studies have shown the role of the anterior STG and the anterior STS in the conceptual representation of social context by the human voice [47–50]. Therefore, our data might suggest that the anterior part of the superior temporal cortex could be recruited to process the social context of human and chimpanzee vocal stimuli. However, this processing would be more automated for the perception of the human voice than for chimpanzee calls because of our high exposure and expertise as humans to these vocal signals, but these hypotheses should be addressed scientifically in studies dedicated to this topic. Our results are also complementary to and coherent with a ‘voice patch’ system in the brain of primates, as put forward by Belin and colleagues [51], and according to which distinct ‘patches’ or subregions of the temporal lobe—especially its anterior portion, would be interconnected and would allow for the processing of voice information. Such system would be present in many primate species such as humans, macaques and marmosets, with most recent evidence suggesting a population of neurons in the anterior STG of the macaque brain selective to human voice [52], as also anticipated in another study on macaques and also in the anterior STG [53]. These fascinating and converging results mirror our present data—with chimpanzee calls triggering responses in the anterior STG/TVA of our human participants—and strongly emphasize the need for pursuing a comparative approach in order to clarify the cross-species neural bases underlying the processing of human and nonhuman primate vocal signals. As we mentioned previously, these interpretations are, considering our results, free of any potential attentional bias towards one species over the others, since no effect was observed on that matter in a control, behavioral study involving an independent sample of twenty-eight participants in a species-specific exogenous cueing attentional paradigm—Methods and Fig.S1.

Importantly, our data also emphasize the influence of acoustic features and especially acoustic proximity between human and chimpanzee vocalizations: we show that activity in the anterior STG and more generally in the anterior TVA partly depends on phylogenetic and importantly on key acoustic features—including acoustic proximity. Consistent with previous studies [27, 28], we did not expect TVA activity for macaque calls processing because they are both phylogenetically and acoustically more distant from humans than the other species in this study—although, as mentioned above, we found a very small cluster in the left primary auditory cortex and mid STS for model 3. It is interesting to note that in model 2, with acoustic distance as trial-level covariate, we observed TVA activity only for chimpanzee but not macaque calls, giving further weight to the importance of acoustic distance in this context. On this matter, the mid STS location of the macaque-related cluster aligns with a cortical region preferentially responsive to non-rhythmic, ‘noise-like’ sounds—a category that acoustically includes macaque vocalizations—rather than to harmonic, voice-like signals [54]. This is further consistent with the posterior-to-anterior functional gradient of the superior temporal lobe, whereby mid and posterior regions process lower-level acoustic features and more anterior STG/STS regions support categorical sound recognition [55]. Also, if only phylogenetic proximity mattered, bonobo calls should also elicit activity in the TVA because they are as phylogenetically close to humans as chimpanzees. But this viewpoint is rather reductive and our results show that this is not entirely correct, and that activity in the TVA crucially depends on the acoustic properties of the perceived vocalizations since we cannot infer phylogeny from vocalizations. This interpretation is strongly supported by the inclusion of acoustic Mahalanobis distance for each species compared to the human voice as a trial-level covariate of no-interest. Using such modelling, differential neuroimaging results between chimpanzee and bonobo vocalizations were explained by both acoustic and phylogenetic proximity in the TVA. These results are consistent with the recent proposal—and recent findings [33]—that there are substantial differences between chimpanzee and bonobo vocalizations. These encompass fundamental frequency range and mean due to larynx length [31, 35]—despite the evolutionary relatedness to chimpanzees [31]. Therefore, the interaction between phylogeny and acoustic distance or proximity would explain the anterior TVA expansion for processing specifically chimpanzee but not bonobo vocalizations. This argument however falls short to explain the activations of the TVA by macaque calls in model 3, that may have been treated as less complex or structured sounds in the mid TVA [54].

Overall, it seems reasonable to hypothesize that TVA activity is not *per se* human-specific [2, 49], but that TVA are instead sensitive to vocalizations from other primate species, provided that these vocalizations have sufficient acoustic proximity to human vocal signals—which would in itself be related to anatomical and/or behavioral changes throughout phylogenetic evolution. This integrative view is again consistent with the concept of a ‘voice patch’ system in the primate brain [51]. We therefore propose that the mid and anterior TVA, unlike the rest of the TVA, would be heterospecific—sensitive to vocalization acoustics triggered by evolution. This proposition also implies a validation of our mechanistic hypothesis according to which the mean fundamental frequency of chimpanzee but not bonobo calls—the former being much closer to the mean fundamental frequency of the human voice [31], would allow for a better identification and recognition of chimpanzee calls by humans. This advantage would rely on neurons of the human auditory cortex—both the primary and more secondary regions—being specialized in the processing of low to mid fundamental frequencies such as those of the human voice and chimpanzee calls. In our third analysis model, we looked further into this aspect and included several acoustic properties of our stimuli as a function of the four species in our stimuli. A discriminant analysis [33] allowed us to select specific acoustic features that best discriminate between our species stimuli. Namely, we took the six parameters explaining the most the differences between our stimuli, including vocalization loudness, intensity—similar to our ‘energy’ covariate of model 1, in addition to change in spectrum, F2 bandwidth contour, F0 power and intensity contour difference. Using these more sophisticated acoustic features as covariates of no-interest, we still obtained brain imaging results very similar to those of model 1 and even closer to model 2—with acoustic distance as covariate, yet with some subtle differences in anterior STG cluster size and location. The peaks were indeed located more ventral and were larger—as compared to results of model 1 & 2, especially for the processing of chimpanzee-selective and macaque-selective vocalizations compared to all primate species and to nonhuman primates alone. These results suggest that the inclusion of spectrum change to intensity- and frequency-related acoustical parameters of the vocal signals slightly shifted and enlarged activation locations in the anterior STG. This result is again congruent with the proposed existence of ‘voice patches’ in the temporal lobe of primate species [51], with the interconnectivity of these patches highly depending on very fine-grained acoustic aspects of primate vocal signals. This step motivated the inclusion of these parameters as covariates of no-interest in neuroimaging model 3, to retain brain activations marginally independent of such acoustics. The congruence between these data should be explored in more detail in the future by the combination of computational bioacoustics and functional neuroimaging, due to the high relevance and sensitivity of combining these techniques to investigate primate social communication [38].

A final but maybe more secondary interpretation arising from our results regarding bonobo calls also supports the evolutionary divergence of this peculiar species. According to the self-domestication hypothesis, bonobos would have evolved differently than chimpanzees due to selection against aggression [56]. Interestingly, differentiation in the evolutionary path of bonobos has influenced both their behavior [34] and morphology, leading to differences at the level of call production [31, 36]. Considering these documented acoustic differences and putting them in perspective with our neuroimaging data, the calls of our last common ancestor with the other Pan species 8 million years ago [57], may have been closer to those uttered by modern chimpanzees than to those of bonobos. Our data indeed show that modern human brains remain more sensitive to the acoustic characteristics of the calls of the former compared to the latter, arguing for more conserved calls between modern chimpanzees and humans. This aspect is also in line with significant differences between, for instance, the fundamental frequency of human baby cries or babbling (∼250-600Hz) compared to that of bonobos (∼1000-3500Hz) [58, 59], while they correspond more closely to the fundamental frequency of chimpanzee calls (∼500-1000Hz) [31]. In our study, bonobo calls definitely are so much different than those uttered by the species of our other stimuli, that they presumably fall outside of the phylogeny and acoustic proximity factors that we outlined so far. This would also put into perspective the selectivity of mid TVAs for macaque calls. The distinct analysis we computed, dedicated to acoustic differences between chimpanzee and bonobo calls, highlighted specificities in pitch dynamics, phonation quality—periodicity, breathiness, glottal irregularity—and vocal tract resonance. This analysis therefore revealed the more stable, lower-pitched, periodically structured nature of the calls of chimpanzees relative to the higher, breathier, and more variable calls of bonobos. These data align well with those obtained through Mahalanobis acoustic distance data discussed above, again converging towards voice-like acoustics for chimpanzee calls.

In a sense, we therefore validate our first hypothesis regarding the existence of acoustic distance between each primate species used in our study. We also partially validate our second hypothesis, albeit not completely. In fact, macaque compared to bonobo or other primate calls in model 3 revealed mid TVA activations, and we think that these activations may depend specifically on the importance of the most discriminant acoustic features. Several TVA subregions or ‘patches’ underlying cross-species primate vocalization processing might therefore exist, and our data highlight at least one of them in the mid and anterior portion of the TVA.

Behavioral data should also be discussed in the context of the species categorization task. We know from our attentional control task that species categorization was not biased by greater saliency from a specific species. We can therefore consider the proportion of correct categorization as attention-bias free in that regard. A ceiling effect was expected and observed for human voice, while only chimpanzee and macaque vocal signals were categorized above chance level for nonhuman primate species. Interestingly, no difference was observed in accuracy between these two, while bonobo calls were categorized below chance. These results are at odds with the acoustic distance argument *per se*, since we show that macaque coos are the furthest from human voice and yet these were better recognized than acoustically-closer bonobo calls. The neural correlates of species categorization performance also point towards the calls of bonobos being completely different than the other species, recruiting most TVA territory compared to other species. Combined, and aligning with previous data [33, 37, 39], these puzzling behavioral and neural results illustrate the very different nature of bonobo calls, and the fact that acoustically, they may have even represented an ‘object’ that was not part of the primate category for our participants. Considering the vast TVA activations compared to the other species, bonobos calls were also most probably very engaging, congruent with massive resources being allocated due to hesitation and decisional difficulty [37].

Considering behavioral data and acoustics, we should also mention the possibility that familiarity to some species may have influenced our results. In fact, differential familiarity with the stimulus species through media or zoo exposure cannot be entirely ruled out. However, this interpretation is undermined by the observation that bonobo calls failed to elicit any anterior TVA response. This interpretation also suffers from the fact that bonobos phylogenetically equidistant from humans that chimpanzees, and are equally unfamiliar to Western participants, although maybe less culturally present than chimpanzees. Consistent with this nuanced viewpoint, ERP evidence suggests that early, automatic neural responses to primate vocalizations are modulated by phylogenetic proximity rather than familiarity—with the latter influencing only later attentional processing stages [60, 61].

A fundamental challenge, shared by vocal science, cognitive and affective neuroscience fields, relates to dissociating biologically relevant information from acoustic properties: species-specific vocal signals are in fact entirely instantiated in acoustics—there is no ‘non-acoustic’ biological residual in the strict sense [33, 38, 55]. Our models (1, 2 & 3) therefore demonstrate independence from specific, theory-motivated parameterizations of the acoustic signal rather than from acoustic information in its entirety. Therefore, residual anterior TVA response to chimpanzee calls most plausibly reflects sensitivity to higher-order acoustic structure not captured—at least not entirely—by these lower-dimensional features. This higher-order acoustic structure may include patterns of spectro-temporal properties of species-typical calls, their sequential organization, or the results could also depend on an evolutionary ‘tuning’ of the human anterior TVA to an acoustic processing niche shared with the great ape vocal system [51]. We acknowledge this as a fundamental limitation of the acoustic-control approach in comparative vocal neuroimaging, and in our study, and suggest that encoding models using richer spectro-temporal representations—such as, for instance, more complex sequences of calls rather than single bursts—may offer more exhaustive acoustic control in future work.

Therefore, even though we tried to control for critical acoustic features, species categories and their related evolutionary distance, several limitations should be mentioned. These limitations are both theoretical and methodological. First, we cannot rule out the fact that including more primate species in our set of stimuli would not have influenced the results. In fact, even though our species categories were specifically chosen for this task, the inclusion of vocalizations from other great apes—such as gorillas or orangutans—would have broadened the scope of our results. Related to this aspect, we can also mention that tackling primate phylogeny, which spans over millions of years, with only four species restricts the possible inference based on our results. Also, not including either nonvocal stimuli or scrambled nonhuman primate calls limits the interpretability of the results, since we cannot rule out that the auditory species ‘object’ per se was triggering the results. Active species categorization compared to passive listening may also introduce a decision-making bias in the neural effects observed. In fact, we observed improved sensitivity of our data by the use of more sophisticated acoustic modeling, namely the inclusion of both between-species acoustic distance and of the most discriminant acoustic features in the functional imaging data. However, we did not include as stimuli—or in a control task—the synthesized acoustic parameters of interest, for instance by using species-specific F0 contour or its spectral content in other neutral, comparable auditory stimuli. We cannot therefore *completely* rule out that such task would not trigger brain activations that overlap with our results—although such data would not be mutually exclusive with our data and interpretation. Future work should therefore address with the greatest level of detail the specific question of acoustics in primate vocalization processing, in addition to adding more—as well as synthesized—stimuli from other great ape species, and if possible using both bursts as well as more complex sequences of calls. The origin of these acoustic differences should also be investigated, since we can assume that these differences originated at least partially from evolutionary processes as well as survival and adaptation mechanisms. Linked to this aspect, phylogeny and acoustics were at least partly confounded in our factorial design, and using dedicated synthetic vs. natural vocal material as stimuli would disentangle this issue. Setting the acoustic covariates to zero in the fMRI models also did not inform on the impact of having acoustics of interest, and this aspect should be studied in the future. Finally, individual differences in the processing and preference of one species over another or over all the others cannot be ruled out, even though we provide evidence that attentional effects toward the vocalizations of a specific species did likely not exist in our data. Therefore, individual differences should be assessed in more detail in the future, with the inclusion of participant-level covariates such as questionnaire scores assessing the familiarity with primate vocalizations or the hedonic value of these vocalizations for each individual. Among the more general limitations of nonhuman primate neuroscience lies the fact that more inclusive and large-scale collaborations would be needed. Such collaborations and framework would lead to a better study and understanding of primate neuroscience, and previous initiatives have recently been put forward in this direction [62, 63].

Taken together, our data suggest that phylogeny-driven specific acoustic features appear to be necessary to trigger cross-species activity in the human temporal voice areas—especially in subregions of the TVA in which increased activity underlies voice signals compared to animal and nature sounds. We provide evidence for specific anterior and mid TVA subregions that underlie the processing of the calls of one of our closest relatives, namely chimpanzees but also of macaques, respectively. In line with recently reported literature, we contend that the human TVA are also involved in the processing of heterospecific primate vocalizations, provided they exhibit sufficient phylogenetic and especially spectro-temporal acoustic proximity to the human voice; as such, we predict that other similarities will be uncovered in the processing of human and nonhuman primate communicative signals. Finally, our results support a critical evolutionary continuity between the structure of human and chimpanzee vocalizations, possibly reflecting one of their common ancestors, as opposed to bonobo vocalizations that underwent more recent and critical changes within the last 1-2 million years. In contrast, the chimpanzee vocal system may be closer to the one of the common ancestors of humans and chimpanzees, as shown by the conserved activation in the human modern brain.

## Material and Methods

### Species categorization task

#### Participants

Twenty-five right-handed, healthy, either native or highly proficient French-speaking participants took part in the study. One participant was excluded because he had no correct response at all and may have fallen asleep, while another participant was excluded due to incomplete scanning and technical issues at the MRI scanner, leaving us with twenty-three participants (10 female, 13 male, mean age 24.65 years, SD 3.66). With this sample size and our study design, we achieved a power of 75.12% for a between-means comparison with Effect size dz=0.5 and alpha=0.05 as calculated in G*Power version 3.1.9.7 [64]. All participants were naive to the experimental design and study, had normal or corrected-to-normal vision, normal hearing and no history of psychiatric or neurologic incidents. Participants gave written informed consent for their participation in accordance with ethical and data security guidelines of the University of Geneva. The study was approved by the Ethics Cantonal Commission for Research of the Canton of Geneva, Switzerland (CCER) and was conducted according to the Declaration of Helsinki.

#### Stimuli

Seventy-two vocalizations of four primate species (human, chimpanzee, bonobo, and rhesus macaque) were used in this study (see Fig.1**A**). The eighteen selected chimpanzee, bonobo and rhesus macaque vocalizations contained single calls or call sequences produced by 6 to 8 different individuals in agonistic (threat, distress) or affiliative (‘positive’) social contexts. These were randomly selected—in a between-participants fashion—among our full database of primate stimuli, containing specifically: 15 chimpanzee individuals (recorded in the wild in the Budongo forest, Uganda), 10 bonobo individuals (recorded in the wild in the Salonga national park, Democratic Republic of Congo, DRC) and 16 macaque individuals (recorded in the wild from semi-free monkeys on Cayo Santiago, Puerto Rico). We then selected eighteen human voices obtained from a nonverbal validated stimuli set of Belin and collaborators [65], which were expressed by two male and two female adults expressing positive or negative social interactions. All vocal stimuli were standardized to 750 milliseconds using PRAAT (www.praat.org) but were not normalized in any way in order to preserve the naturalness of the sounds [40] and to allow for low-level acoustic parameters to be used in neuroimaging data modelling.

#### Experimental procedure and paradigm

Laying comfortably in a 3T scanner, participants listened to a total of seventy-two stimuli pseudo-randomized and played binaurally using MRI compatible earphones at 70 dB SPL (Model ‘S14’, Sensimetrics Corporation, Gloucester, MA, USA). At the beginning of the experiment, participants were instructed to identify the species that expressed the vocalizations using a keyboard. Three vocalizations of each species were first presented outside the scanner to the participants, revealing to them which species the stimuli were from—as a form of familiarization to the auditory material and the species. These stimuli were then discarded from the task. For instance, the instructions could be “Human – press 1, Chimpanzee – press 2, Bonobo – press 3 or Macaque – press 4”. The pressed keys were pseudo-randomly assigned across participants (response box: fORP, Cortech Solutions, Inc., Wilmington, NC, USA). In a 3-5 second interval (jittering of 400 ms) after each stimulus, participants were asked to categorize the species. If the participant did not respond during this interval, the next stimulus followed automatically. See Fig.1**A** for a detailed illustration of the paradigm.

### Temporal voice areas localizer task

#### Participants

Two independent samples of participants performed the task while undergoing fMRI scanning: the sample of this study (10 female, 13 male, mean age 24.65 years, SD 3.66), leading to the delineation of sample-specific TVA; an independent sample of ninety-eight right-handed, healthy, either native or highly proficient French-speaking participants (52 female, 46 male, mean age 24.66 years, SD 4.97) leading to the delineation of more general and representative TVA. All participants were naive to the experimental design and study, had normal or corrected-to-normal vision, normal hearing and no history of psychiatric or neurologic incidents. Participants gave written informed consent for their participation in accordance with ethical and data security guidelines of the University of Geneva. The study was approved by the Ethics Cantonal Commission for Research of the Canton of Geneva, Switzerland (CCER) and was conducted according to the current regulations in Switzerland.

#### Stimuli and paradigm

Auditory stimuli consisted of sounds from a variety of sources [2]. Vocal stimuli were obtained from 47 speakers: 7 babies, 12 adults, 23 children and 5 older adults. Stimuli included 20 blocks of vocal sounds and 20 blocks of non-vocal sounds. Vocal stimuli within a block could be either speech 33%: words, non-words, foreign language or non-speech 67%: laughs, sighs, various onomatopoeia. Non-vocal stimuli consisted of natural sounds 14%: wind, streams, animals, 29%: cries, gallops, the human environment, 37%: cars, telephones, airplanes or musical instruments, 20%: bells, harp, instrumental orchestra. The paradigm, design and stimuli were obtained through the Voice Neurocognition Laboratory website (http://vnl.psy.gla.ac.uk/resources.php). Stimuli were presented through earphones (Model ‘S14’, Sensimetrics Corporation, Gloucester, MA, USA) at an intensity that was kept constant throughout the experiment 70 dB sound-pressure level. Participants were instructed to actively listen to the sounds. The silent inter-block interval was 8 s long.

### Exogenous attention task using species as cues

This task was performed to control for potential attentional biases or specific salience of one species compared to the others. This task was therefore a control task and the results are reported in Fig.1. The task was designed according to work on attention orienting following vocal material presentation [66]. More specifically, a cue is first presented and is quickly followed by the presentation of a neutral target to detect. The detection of this target can reliably be delayed or accelerated depending on the nature of the cue. In this control study, the cue corresponded to the stimuli of the main species categorization task followed by a sine wave tone (a ‘bip’) that had to be detected as fast as possible (the target). Any specific attentional bias of a species would therefore trigger reaction times differences for target detection, while no differences would invalidate any attentional effect or increased salience linked to a specific species. We did not observe any difference between cue species in this task (χ^2^(2)=3.33, *p*=0.34, Fig.1).

#### Participants

Twenty-eight participants (independent of the samples presented so far) took part in this behavioral study (15 female, 13 male, mean age 22.63 years, SD 5.00). With this sample size and our study design, we achieved a power of 82.48% for a between-means comparison with Effect size dz=0.5 and alpha=0.05 as calculated in G*Power version 3.1.9.7 [64]. All participants were naive to the experimental design and study, had normal or corrected-to-normal vision, normal hearing and no history of psychiatric or neurologic incidents. Participants gave written informed consent for their participation in accordance with ethical and data security guidelines of the University of Geneva. The study was approved by the Ethics committee of the University of Geneva (Switzerland), Department of Psychology and Educational Sciences, and was conducted according to the Declaration of Helsinki.

#### Stimuli and paradigm

The cues of the task were the species stimuli (human voice, chimpanzee, bonobo and macaque calls) used in the species categorization task above. The target ‘bip’ was a sine wave tone created using Matlab (Matlab 2020a, The Mathworks, Inc., Natick, MA, USA) with a wave frequency of 600Hz, a fade-in and fade-out of 10 ms each and a total duration of 100 ms. The auditory material was presented through headphones (Sennheiser HD-25 II, Sennheiser electronic SE & Co. KG, Germany) at a constant sound-pressure level of 70dB. The procedure unfolded as follows, on a computer with a light grey screen background: for each trial (N=172), a cue (human voice, chimpanzee, bonobo or macaque call) was presented for 750 ms while a black fixation cross was presented at the center of the screen. Following a jittered blank screen of duration 100 to 250 ms (in steps of 50 ms), the target bip was presented for 100 ms. Right at the end of the presented of the bip, the fixation cross turned white, indicating that the response screen had started and that a response was expected as fast and accurately as possible as instructed, by using the ‘space’ key of the keyboard. A varying inter-trial interval of 1 s to 2.5 s was used (in steps of 500 ms). Among the total of 172 trials, there were 24 trials per species (12 in agonistic and 12 in affiliative social contexts) for a total of 96 stimuli (4 species * 24 trials=96) that were each presented twice (N=96*2=172).

### Behavioral data analysis

#### Species categorization task

##### Accuracy

Behavioral data were used to exclude participants who had below chance level categorization of human voices. Therefore, data from twenty-three participants mentioned in the *Species Categorization Task - Participants* section above were analyzed using R studio software (R Studio team [67] Inc., Boston, MA, url: http://www.rstudio.com/) and the lme4 [68] and emmeans [69] packages. These data can be found in a published article focused on decisional aspects in the frontal cortex—region-of-interest analysis and computational modelling of the probability of correct species categorization—using the same species stimuli as in this study, but with different modeling of acoustics [37].

For the present study, behavioral data were analyzed in a similar fashion as previously published by Ceravolo et al. [37] but with acoustic covariates of Neuroimaging model 3, as described below (vocalization loudness, intensity, change in spectrum, bandwidth contour of the second formant, power of the fundamental frequency and the difference in intensity contour). Namely, the dependent variable was a binarized accuracy for each trial (*response*), determining whether the participants categorized the species correctly or incorrectly (1 or 0, respectively). A logistic regression was used to predict this binary response variable with interacting *Species* and vocalization *Context*, in addition to the main effect of each of the 6 most discriminant acoustic parameters as fixed effects. Random effects included the identity of the participants (*ParticipantID*), response reaction times (*RT*)—as these were of no interest but worth controlling for, participant gender (*Gender*) and age (*Age*), and stimulus identity (*StimID*). This model explained 62.10% of the variance in the data (R2c).

The formula, subsequently referred to as Neuroimaging model 4, was the following:

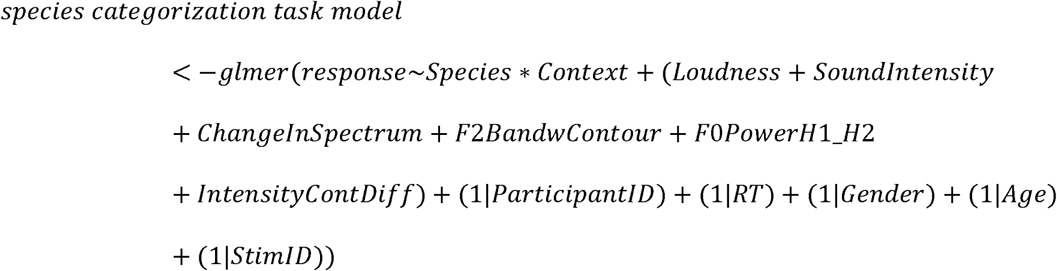

#### Exogenous attention task

##### Reaction times

In this study, the dependent variable of interest was the reactions times to detect the target ‘bip’. In order to remove extreme values, we discarded for each participant the values below the 5^th^ percentile and above the 95^th^ percentile. In average, the number of trials per participants therefore went down from 172 to 165 (∼4.1% of trials removed). Data were then analyzed using R studio software (R Studio team [67] Inc., Boston, MA, url: http://www.rstudio.com/) using linear mixed effects modeling of the lme4 package [68]. The formula was the following:

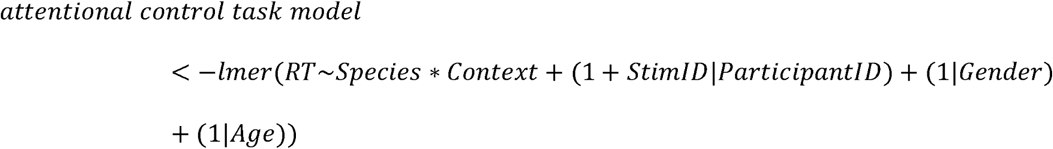

in which *RT* is the reaction times (dependent variable), *Species* is the four species of the stimuli (human, chimpanzee, bonobo, macaque) interacting with the *Context* of production (affiliative, agonistic) (fixed effects). The random effects were: the random slope of the identity of each stimulus (*StimID*) as a function of participants (*ParticipantID*), participant gender (*Gender*) and age (*Age*). This model explained 57.54% of the variance in the data (R2c).

There was no significant effect of Species (χ^2^(3)=3.33, *p*=.34), a significant effect of Context χ^2^ (2)=11.39, *p*<.01) and no interaction between Species and Context (χ^2^(6)=3.06, p=.80). The effect of Context was explained by slower reaction times for affiliative than agonistic vocalizations, independent of Species (χ^2^(1)=6.08, *p*<.05). Descriptive statistics per Species for the reaction times were the following: Human, mean=207.28, SD=118.43; Chimpanzee, mean=211.31, SD=127.07; Bonobo, mean=209.45, SD=151.85; Macaque, mean=214.05, SD=117.84. Descriptive statistics per Context for the reaction times were: Affiliative, mean=215.03, SD=115.41; Agonistic, mean=208.16, SD=135.39. See illustration in Fig.S1.

### Acoustic analysis of the vocalizations

#### Mahalanobis acoustic distance (Neuroimaging model 2)

To quantify the impact of acoustic similarities in human recognition of affective vocalizations of other primates, we extracted 88 acoustic parameters from all vocalizations using the extended Geneva Acoustic parameters set defined as the optimal acoustic indicators related to voice analysis (GeMAPS [42]). This open-source set of acoustical parameters was selected based on: i) their potential to index affective physiological changes in voice production, ii) their proven value in former studies as well as their automatic extractability, and iii) their theoretical significance. GeMAPS relies on an automatic extraction system, which therefore automatically extracts an acoustic parameter set from an audio file—in an unsupervised, minimalistic manner. Then, to assess the acoustic distance between vocalizations of all species, we ran a General Discriminant Analysis model (GDA). More precisely, we used the 88 acoustical parameters in a GDA in order to discriminate our stimuli based on the different species (human, chimpanzee, bonobo, and rhesus macaque). Among these 88 acoustical parameters, we excluded those that were strongly correlating—i.e., with correlation scores r>.70—to avoid redundancy and minimize multicollinearity. Following this selection process and the GDA, we eventually retained 16 acoustic parameters (Table S1). Namely, the parameters were, in that order of importance: 1) Arithmetic mean of the contour of the melodic-frequency cepstral coefficient 4; 2) Arithmetic mean of the contour of the frequency with most flux around it (change in spectrum); 3) Normalized standard deviation of the contour of the frequency with most flux around it (change in spectrum); 4) Conversion of power to decibel of the root mean square of the sound energy; 5) Arithmetic mean of the slope of timeframe 0-500; 6) Arithmetic mean of the contour of the melodic-frequency cepstral coefficient 1; 7) Arithmetic mean of the second formant amplitude power log relative to fundamental frequency; 8) Normalized standard deviation of the contour of the second formant amplitude power log relative to fundamental frequency; 9) Arithmetic mean of power log relative to fundamental frequency of the harmonic one and two; 10) Normalized standard deviation of the contour of the relative amplitude difference in local dB; 11) Arithmetic mean of the contour of the sum of the auditory spectrum; 12) Rate of peaks per second of the sum of the auditory spectrum; 13) Normalized standard deviation of the contour of the first formant amplitude power log relative to fundamental frequency; 14) Arithmetic mean of the contour of the bandwidth of the second formant; 15) Arithmetic mean of the contour of the relative amplitude difference in local dB; 16) Normalized standard deviation of the contour of the melodic-frequency cepstral coefficient 2.

We subsequently computed multidimensional Mahalanobis distances to classify the 72 stimuli on these selected acoustical features. A Mahalanobis distance is a generalized pattern analysis comparing the distance of each vocalization from the centroids of the different species vocalizations. This analysis allowed us to obtain an acoustical distance matrix used to test how the acoustical distances were differentially related to the different species (see Fig.1**B**) and we used it as a covariate of no-interest in neuroimaging model 2. Using a one-way ANOVA with the Distance as the dependent variables and the Species as independent variable, the main effect of Species was significant (F(3,88)=15.84, *p*<.001). All between-species differences were significant (.01<*p*<.001; see Fig.1 and Table S2).

These data are the topic of a publication about the impact of acoustic parameters on the recognition of the affective cues of primate vocalizations by human participants [33]. All details are described in this article for the present stimuli.

#### Most discriminant low-level acoustic parameters (Neuroimaging model 3)

Following the GDA on the 88 acoustic parameters [42, 43] of the species stimuli presented above, we decided to use as covariates of no-interest the most discriminant low-level acoustic features of our stimuli to maximize brain activations that are independent of these features. We therefore included the most significant acoustic features ([r>.70] and [r<-.70]) of the first three factors of the GDA, that explained 27.14%, 21.63% and 18.99% of the variance, respectively [33]. Such selection left us with the following acoustic features: Factor 1, (1) vocalization loudness (r=0.92), (2) intensity (r=0.87), (3) change in spectrum (r=0.72); Factor 2, (4) bandwidth contour of the second formant (F2; r=0.79); Factor 3, (5) power of the fundamental frequency (F0; r=0.80) and finally (6) the difference in intensity contour (r=-0.71). The acoustic parameters were used as covariates of no-interest (N=6) in that specific order—namely, from the highest to lowest factor saturation—in neuroimaging model 3.

Again, all details of this analysis are described in detail in a dedicated article for the present stimuli [33] and the values are reported in Table S1.

#### Acoustic parameters characterizing chimpanzee and bonobo calls

We performed a principal component analysis (PCA) on the 88 GeMAPS [42, 43] acoustic parameters by using only the stimuli recorded from chimpanzee and bonobo individuals, in order to constrain the extraction of acoustic features of interest to these two species only. We again included the most significant acoustic features and took care of multicollinearity (excluding acoustic parameters highly correlating together with [r>.70] and [r<-.70]) before PCA. This analysis left us with 13 principal components (PC) accounting in total for at least 80% of the variance. Variance explained specifically for each PC (and its name and loading) was: PC1 (F0 Range (semitones), 0.349), 12.47%, PC2 (Spectral Slope 500–1500 Hz Variability, +0.441), 9.50%; PC3 (F1 Frequency Variability, +0.357), 8.87%; PC4 (Unvoiced Segment Duration SD, +0.432), 7.38; PC5 (HNR Variability, +0.339), 6.83%; PC6 (Spectral Slope 0–500 Hz (mean), +0.474), 6.08%; PC7 (F1 Bandwidth Variability, +0.414), 5.81%; PC8 (F0 Rising Slope Variability, -0.403), 5.22%; PC9 (F2 Amplitude Variability (relative to F0), +0.387), 5.09%; PC10 (Spectral Slope 0–500 Hz Variability, +0.358), 4.38%; PC11 (F3 Amplitude Variability (relative to F0), +0.520), 4.10%; PC12 (Jitter Variability, +0.407), 3.77%; PC13 (H1–A3 Amplitude Ratio Variability, +0.470), 3.67%. Cumulative variance among the 13 PC was 83.16%.

### Imaging data acquisition

#### Species categorization task

Structural and functional brain imaging data were acquired by using a 3T scanner Siemens Trio, Erlangen, Germany with a 32-channel coil. A 3D GR\IR magnetization-prepared rapid acquisition gradient echo sequence was used to acquire high-resolution (0.35 x 0.35 x 0.7 mm^3^) T1-weighted structural images (TR = 2400 ms, TE = 2.29 ms). Functional images were acquired by using fast fMRI, with a multislice echo planar imaging sequence with 79 transversal slices in descending order, slice thickness 3 mm, TR = 650 ms, TE = 30 ms, field of view = 205 x 205 mm2, 64 x 64 matrix, flip angle = 50 degrees, bandwidth 1562 Hz/Px. In total for this task, 636 functional volumes of 79 slices were acquired for each participant for a total of 50244 slices per participant. For our whole sample of twenty-three participants, 14628 volumes were acquired for a grand total of 1’155’612 slices.

#### Temporal voice areas localizer task

Structural and functional brain imaging data were acquired by using a 3T scanner Siemens Trio, Erlangen, Germany with a 32-channel coil. A magnetization-prepared rapid acquisition gradient echo sequence was used to acquire high-resolution (1 x 1 x 1 mm^3^) T1-weighted structural images TR = 1,900 ms, TE = 2.27 ms, TI = 900 ms. Functional images were acquired by using a multislice echo planar imaging sequence with 36 transversal slices in descending order, slice thickness 3.2 mm, TR = 2,100 ms, TE = 30 ms, field of view = 205 x 205 mm2, 64 x 64 matrix, flip angle = 90°, bandwidth 1562 Hz/Px. In total for this task, 230 functional volumes of 36 slices were acquired for each participant for a total of 8280 slices per participant. For our sample of ninety-eight participants, 22’540 volumes were acquired for a grand total of 811’440 slices. For the sample-specific data (N=23), 5290 volumes were acquired for a grand total of 190’440 slices.

### Wholebrain data analysis

#### Species categorization task analysis within the temporal voice areas

Functional images were analyzed with Statistical Parametric Mapping software (SPM12, Wellcome Trust Centre for Neuroimaging, London, UK). Preprocessing steps included realignment to the first volume of the time series, slice timing, normalization into the Montreal Neurological Institute [70] (MNI) space using the DARTEL toolbox [71] and spatial smoothing with an isotropic Gaussian filter of 8 mm full width at half maximum. To remove low-frequency components, we used a high-pass filter with a cutoff frequency of 1/128Hz. Three general linear models were used to compute first-level statistics, in which each event was modeled by using a boxcar function and was convolved with the hemodynamic response function, time-locked to the onset of each stimulus. In model 1, separate regressors were created for all trials of each species (Species factor: human, chimpanzee, bonobo, macaque vocalizations) and two covariates of no-interest each (mean fundamental frequency and mean energy of each species) for a total of 12 regressors. Finally, six motion parameters were included as regressors of no interest to account for movement in the data and our design matrix therefore included a total of 18 columns plus the constant term. The species regressors were used to compute simple contrasts for each participant, leading to separate main effects of human, chimpanzee, bonobo, and macaque vocalizations. Covariates were set to zero in order to model them as no-interest regressors. In model 2, separate regressors were created for all trials of each species (Species factor: human, chimpanzee, bonobo, macaque vocalizations) and one covariate of no-interest for each species (acoustic distance for each species relative to human voice stimuli) for a total of 8 regressors. Six motion parameters were included as regressors of no interest to account for movement in the data and our design matrix therefore included a total of 14 columns plus the constant term. The species regressors were used to compute simple contrasts for each participant, leading to separate main effects of human, chimpanzee, bonobo, and macaque vocalizations excluding acoustic distance (the covariate was set to zero in order to model it as ‘of no-interest’). In model 3, separate regressors were created for all trials of each species (Species factor: human, chimpanzee, bonobo, macaque vocalizations) and six covariates of no-interest each (vocalization loudness, intensity, change in spectrum, bandwidth contour of the second formant (F2), power of the fundamental frequency (F0) and finally the difference in intensity contour) for a total of 28 regressors. Six motion parameters were included as regressors of no-interest to account for movement in the data and our design matrix therefore included a total of 34 columns plus the constant term. The species regressors were used to compute simple contrasts for each participant, leading to separate main effects of human, chimpanzee, bonobo, and macaque vocalizations. Covariates were set to zero in order to model them as no-interest regressors. Finally, for Model 4 we used a model-based approach in which we modelled behavioral data as covariate for each trial as the main regressor of interest. Specifically, we computed a modeling of the behavioral data in R—see Species categorization task model above in the section on behavioral data analysis. On the whole sample (N=23), we computed a mixed-effects logistic regression in which we predicted response accuracy as a function of the interaction between vocalization Species and Context of production. The six most discriminant acoustic parameters (those of Model 3) were also included as fixed effects, and we controlled for trial-wise reaction times (random effect). The obtained regression values for each trial and participant were then fitted on the whole sample and used as trial-level covariate in the present model, representing the probability of correctly categorizing human, chimpanzee, bonobo and macaque stimuli in our task. The design matrix for each participant therefore included two regressors: (1) each trial onset and (2) the associated behavioral response fitted regression value. The advantage of this approach is that it includes all responses and all trials, including correct and incorrect responses, therefore providing a clear and representative picture of the brain mechanisms underlying species categorization in our task—without losing power. Each model was independent from the others and was computed by rerunning the analysis based on each of the three design matrices, taking as data the final preprocessed data for each participant.

For each first-level model of analysis Models 1-3, each of their respective four simple contrasts were then taken to two flexible factorial second-level analyses. For all of these second-level analyses there were two factors: Participants factor (independence set to yes, variance set to unequal) and the Species factor (independence set to no, variance set to unequal). For Model 4, the second-level modeling was a one-sample t-test of the parametric modulator, namely the fitted regression values of the probability of correctly categorizing the species stimuli—computed on the whole sample of 23 participants. Variance was also set to unequal in this model. For these analyses and to be consistent, we only included participants who were above chance level (25%) in the species categorization task (N=23).

Brain region labelling was defined using xjView toolbox (http://www.alivelearn.net/xjview) implementing the Automated anatomical labelling (‘aal3’) atlas [72]. All neuroimaging activations were thresholded in SPM12 by using a wholebrain (Models 1-3) or TVA inclusive masking (Model 4) voxelwise false discovery rate (FDR) correction at *p*<.05 and an arbitrary cluster extent of k>10 voxels to remove very small clusters of activity. For Models 1-3, the TVA outlined in the figures are therefore only visual outlines of these regions, but no region-of-interest (ROI) analysis was performed here in order to maximize data representativeness—which is impacted negatively by ROI analysis.

#### Temporal voice areas localizer task

Functional images were analyzed with Statistical Parametric Mapping software (SPM12, Wellcome Trust Centre for Neuroimaging, London, UK). Preprocessing steps included realignment to the first volume of the time series, slice timing, normalization into the Montreal Neurological Institute [70] (MNI) space using the DARTEL toolbox [71] and spatial smoothing with an isotropic Gaussian filter of 8 mm full width at half maximum. To remove low-frequency components, we used a high-pass filter with a cutoff frequency of 1/128Hz. A general linear model was used to compute first-level statistics, in which each block was modeled by using a block function and was convolved with the hemodynamic response function, time-locked to the onset of each block. Separate regressors were created for each condition (vocal and non-vocal; condition factor). Finally, six motion parameters were included as regressors of no interest to account for movement in the data. The condition regressors were used to compute simple contrasts for each participant, leading to a main effect of vocal and non-vocal at the first-level of analysis: [1 0] for vocal, [0 1] for non-vocal. These simple contrasts were then taken to a flexible factorial second-level analysis in which there were two factors: Participants factor (independence set to yes, variance set to unequal) and the Condition factor (independence set to no, variance set to unequal). An identical analysis architecture was used to delineate more specific subregions of the TVA according to the type of non-vocal material, with categories: animal sounds, music, nature sounds, artificial noise sounds. Timing onsets of these newly created “events”—including their duration—within each non-vocal block of the task were determined and each main effect contrast computed at the first-level was then taken to a second-level flexible factorial analysis with settings identical to the above. All neuroimaging activations were thresholded in SPM12 by using a voxelwise false discovery rate (FDR) correction at *p*<.05 and an arbitrary cluster extent of k>10 voxels to remove very small clusters of activity. Activation outline for vocal > non-vocal—the contrast revealing the TVA—was precisely delineated for the N=98 and the N=23 samples and overlaid on brain displays of the species categorization task (see Fig.S7). TVA subregions according to the auditory material (specific non-vocal condition blocks material) are reported in Fig.S6.

## Acknowledgements

We thank the Swiss National Science foundation (SNSF) for supporting this interdisciplinary project (grant CR13I1_162720 / 1 to DG-TG), the National Centre of Competence in Research (NCCR) (51NF40-104897 to DG) hosted by the Swiss Center for Affective Sciences, as well as the Fondation Ernst et Lucie Schmidheiny supporting CD. TG was additionally supported by a grant of the SNSF during the final editing of this article (grant PCEFP1_186832). We also thank Katie Slocombe and Zanna Clay for providing the nonhuman primates auditory stimuli and Daphne Bavelier for her advice on the design of the species categorization task. We would like also to acknowledge the staff of the Brain and Behavior Laboratory at the University of Geneva where all data were acquired.

## Author contributions

LC designed the task, acquired part of the data, analyzed the data, designed the figures, wrote and edited the manuscript. CD designed the task, acquired the data, analyzed the data, wrote and edited parts of the manuscript. TG provided theoretical background and edited the manuscript. DG helped design the task, the analyses and edited the manuscript.

## Data availability statement

All data, stimuli and codes used in this article are available in the FAIR-compliant open repository YARETA (URL: https://doi.org/10/g8wkrb).

