## Supplementary material for "Sensitivity of the human temporal voice areas to nonhuman primate vocalizations"

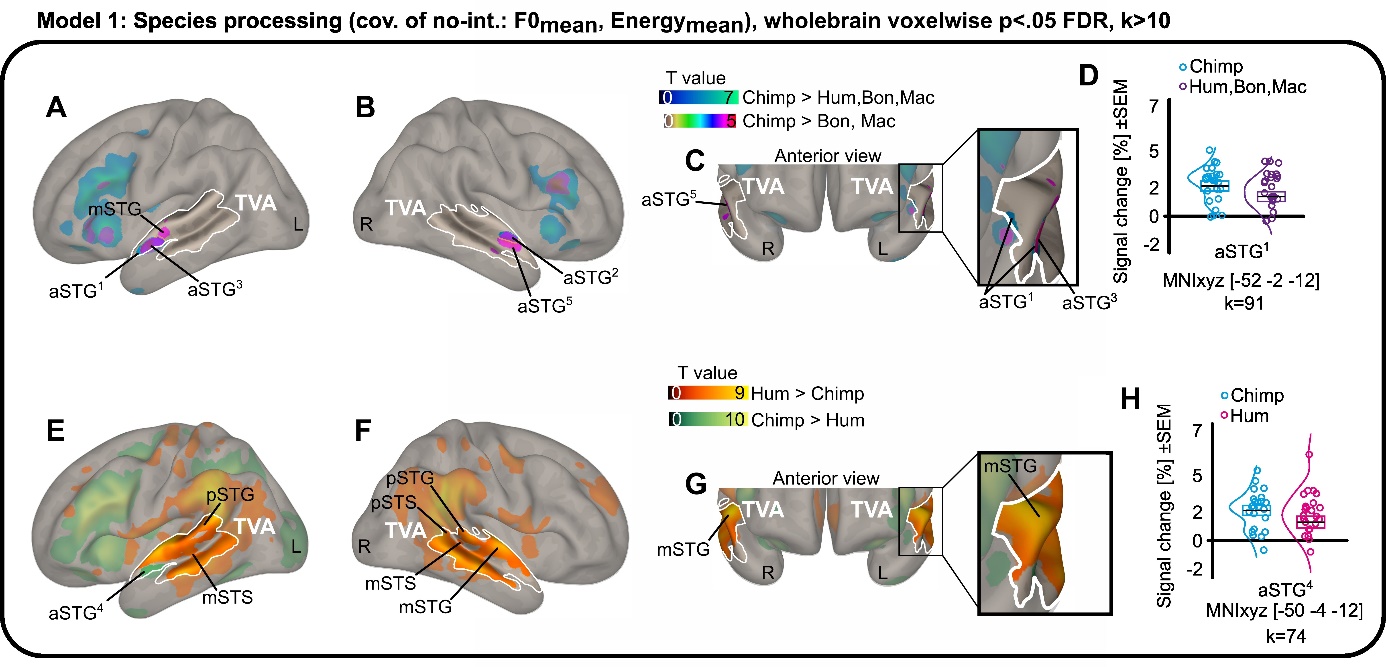

**Fig.S1: Wholebrain results when contrasting the processing of chimpanzee to other species’ vocalizations with mean fundamental frequency and energy as trial-level covariates of no-interest (model 1).** (**A,B,C**) Enhanced brain activity for human and chimpanzee compared to bonobo and macaque vocalizations (purple to yellow) on a sagittal view, overlaid with activity specific to chimpanzee vocalizations (dark blue to green). (**D**) Percentage of signal change for each individual and relevant species according to the contrast in the left anterior superior temporal gyrus (aSTG^1^). Box plots represent mean value (black line) and the standard error of the mean with distribution fit. (**E,F,G**) Direct comparison between human and chimpanzee vocalizations (human > chimpanzee: dark red to yellow; chimpanzee > human: dark green to yellow) as well as between chimpanzee calls vs bonobo and macaque calls (chimpanzee > bonobo and macaque: brown to red) on a sagittal render. (**H**) Percentage of signal change in the anterior superior temporal gyrus (aSTG^2^) when contrasting chimpanzee to human vocalizations for each individual and relevant species according to the contrast with box plots representing mean value (black line) and the standard error of the mean with distribution fit. Brain activations are independent of low-level acoustic parameters for all species (mean fundamental frequency ‘F0’ and mean energy of vocalizations). Data corrected for multiple comparisons using wholebrain voxelwise false discovery rate (FDR) at a threshold of *p*<.05. Percentage of signal change extracted at cluster peak including 9 surrounding voxels, selecting among these the ones explaining at least 85% of the variance using singular value decomposition. Circles represent individual values, boxplot represents the mean and its standard error, and half-violin plots show data distribution. Hum: human; Chimp: chimpanzee; Bon: bonobo; Mac: macaque. TVA: temporal voice areas of an independent sample of N=98. ‘a’ prefix: anterior; ‘m’ prefix: mid; ‘p’ prefix: posterior; STG: superior temporal gyrus; STS: superior temporal sulcus; L: left hemisphere; R: right hemisphere.

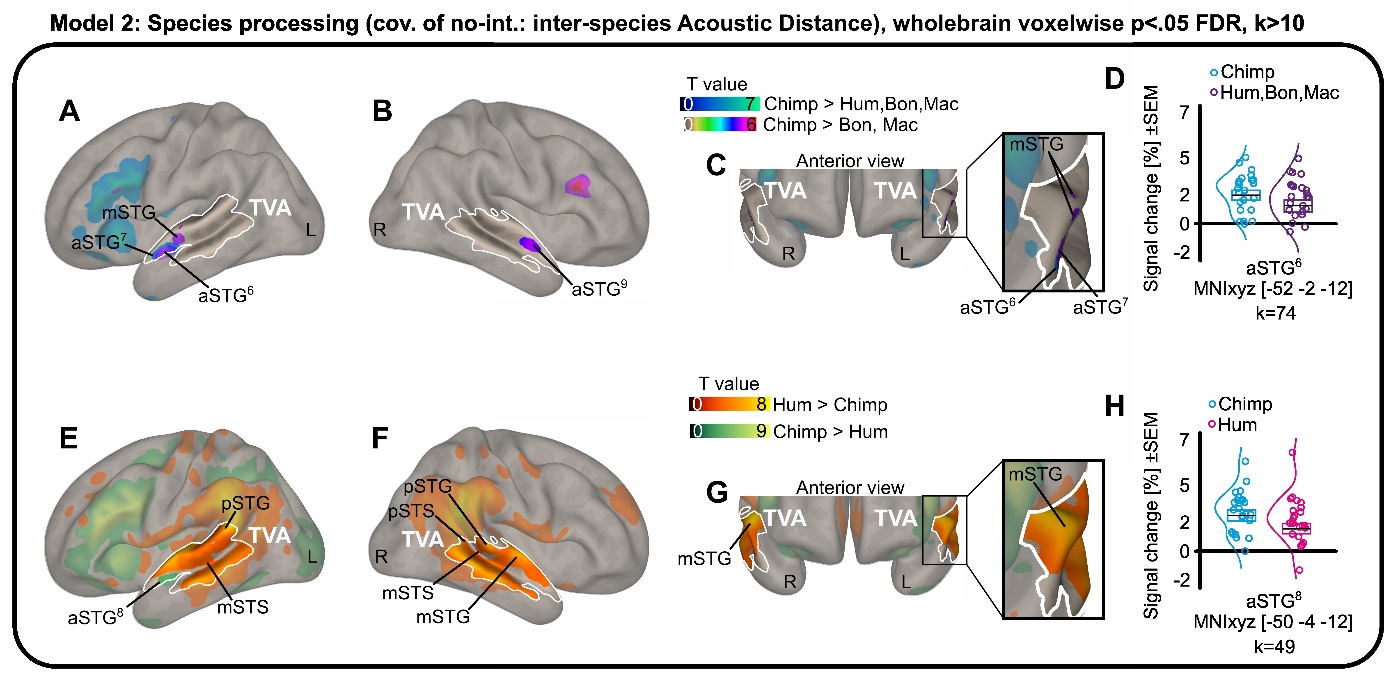

**Fig.S2: Wholebrain results when contrasting the processing of chimpanzee to other species’ vocalizations with Mahalanobis acoustic distance as trial-level covariates of interest (model 2).** (**A,B,C**) Enhanced brain activity for human and chimpanzee compared to bonobo and macaque vocalizations (purple to yellow) on a sagittal view, overlaid with activity specific to chimpanzee vocalizations (dark blue to green). (**D**) Percentage of signal change for each individual and relevant species according to the contrast in the left anterior superior temporal gyrus (aSTG^6^). Box plots represent mean value (black line) and the standard error of the mean with distribution fit. (**E,F,G**) Direct comparison between human and chimpanzee vocalizations (human > chimpanzee: dark red to yellow; chimpanzee > human: dark green to yellow) as well as between chimpanzee calls vs bonobo and macaque calls (chimpanzee > bonobo and macaque: brown to red) on a sagittal render. (**H**) Percentage of signal change in the anterior superior temporal gyrus (aSTG^8^) when contrasting chimpanzee to human vocalizations for each individual and relevant species according to the contrast with box plots representing mean value (black line) and the standard error of the mean with distribution fit. Brain activations are covarying with the acoustic distance of each stimulus for all species. Data corrected for multiple comparisons using wholebrain voxelwise false discovery rate (FDR) at a threshold of *p*<.05. Percentage of signal change extracted at cluster peak including 9 surrounding voxels, selecting among these the ones explaining at least 85% of the variance using singular value decomposition. Circles represent individual values, boxplot represents the mean and its standard error, and half-violin plots show data distribution. Hum: human; Chimp: chimpanzee; Bon: bonobo; Mac: macaque. TVA: temporal voice areas of an independent sample of N=98. ‘a’ prefix: anterior; ‘m’ prefix: mid; ‘p’ prefix: posterior; STG: superior temporal gyrus; STS: superior temporal sulcus; L: left hemisphere; R: right hemisphere.

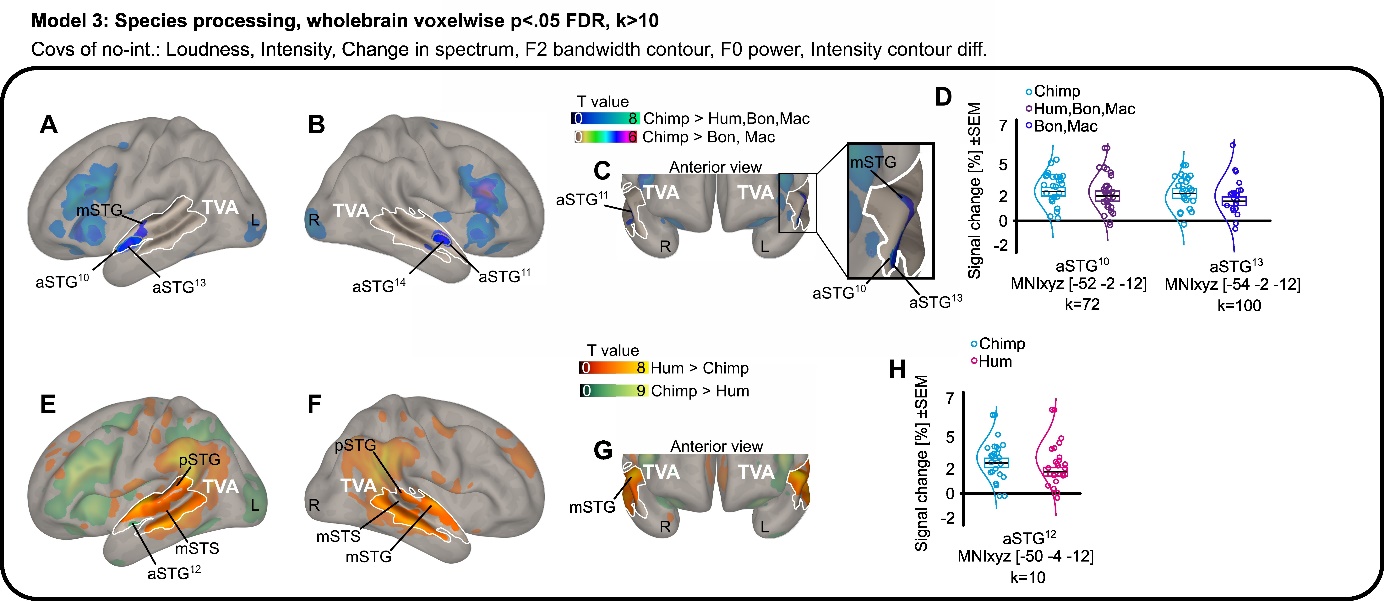

**Fig.S3: Wholebrain results when contrasting the processing of chimpanzee to other species’ vocalizations with vocalization loudness, intensity, change in spectrum, F2 bandwidth contour, F0 power and intensity contour difference as trial-level covariates of no-interest (model 3).** (**A,B,C**) Enhanced brain activity for human and chimpanzee compared to bonobo and macaque vocalizations (purple to yellow) on a sagittal view, overlaid with activity specific to chimpanzee vocalizations (dark blue to green). (**D**) Percentage of signal change for each individual and relevant species according to the contrast in the left anterior superior temporal gyrus (aSTG^10^). Box plots represent mean value (black line) and the standard error of the mean with distribution fit. (**E,F,G**) Direct comparison between human and chimpanzee vocalizations (human > chimpanzee: dark red to yellow; chimpanzee > human: dark green to yellow) as well as between chimpanzee calls vs bonobo and macaque calls (chimpanzee > bonobo and macaque: brown to red) on a sagittal render. (**H**) Percentage of signal change in the anterior superior temporal gyrus (aSTG^12^) when contrasting chimpanzee to human vocalizations and when contrasting chimpanzee to bonobo and macaque calls (aSTG^13^) for each individual and relevant species according to the contrast with box plots representing mean value (black line) and the standard error of the mean with distribution fit. Brain activations are independent of the most discriminant low-level acoustic parameters of the stimuli set. Data corrected for multiple comparisons using wholebrain voxelwise false discovery rate (FDR) at a threshold of *p*<.05. Percentage of signal change extracted at cluster peak including 9 surrounding voxels, selecting among these the ones explaining at least 85% of the variance using singular value decomposition. Circles represent individual values, boxplot represents the mean and its standard error, and half-violin plots show data distribution. Hum: human; Chimp: chimpanzee; Bon: bonobo; Mac: macaque. TVA: temporal voice areas of an independent sample of N=98. ‘a’ prefix: anterior; ‘m’ prefix: mid; ‘p’ prefix: posterior; STG: superior temporal gyrus; STS: superior temporal sulcus; L: left hemisphere; R: right hemisphere.

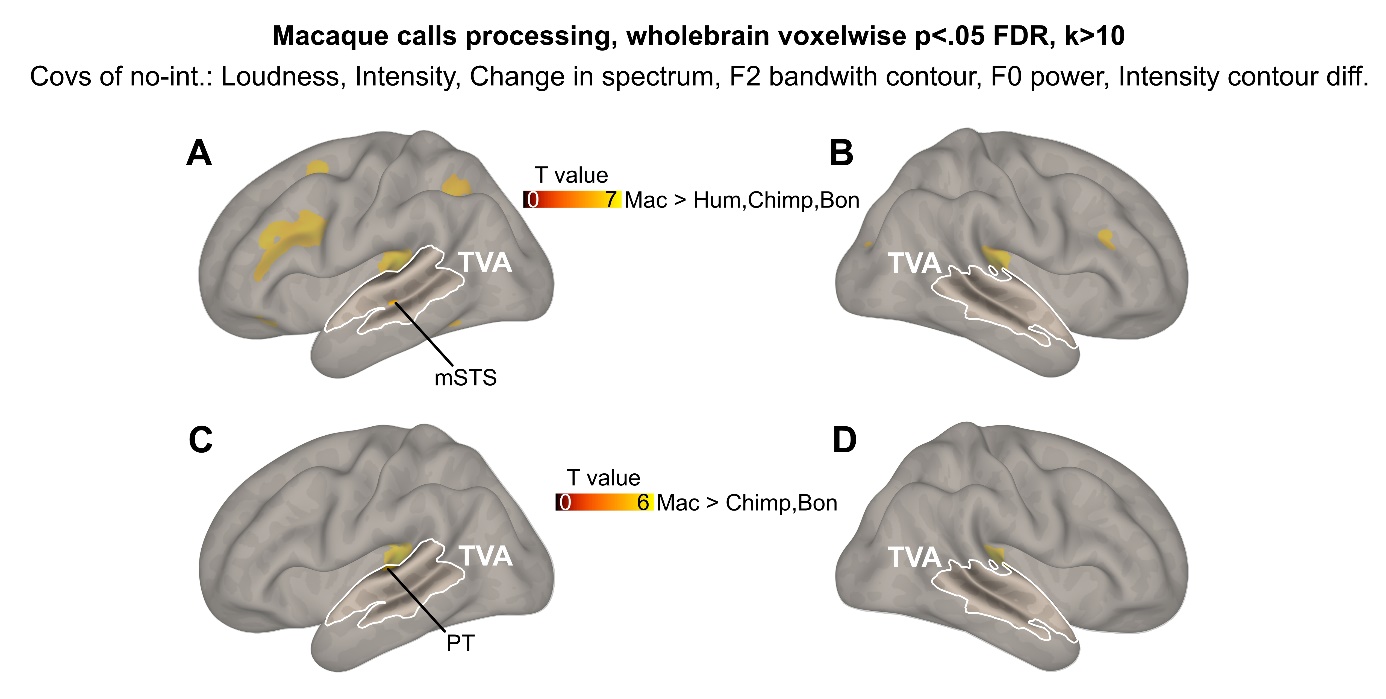

**Fig.S4: Wholebrain activations specific to macaque calls for for model 3.** (**A,B**) Enhanced wholebrain activity for macaque compared to human, chimpanzee and bonobo vocalizations. (**C,D**) Enhanced wholebrain activity for macaque compared to chimpanzee and bonobo vocalizations. Data corrected for multiple comparison using wholebrain voxelwise false discovery rate (FDR) at a threshold of *p*<.05, k=10. Hum: human; Chimp: chimpanzee; Bon: bonobo; Mac: macaque. TVA: sample-specific (N=23) temporal voice areas; PT: planum temporale; mSTG: mid superior temporal gyrus.

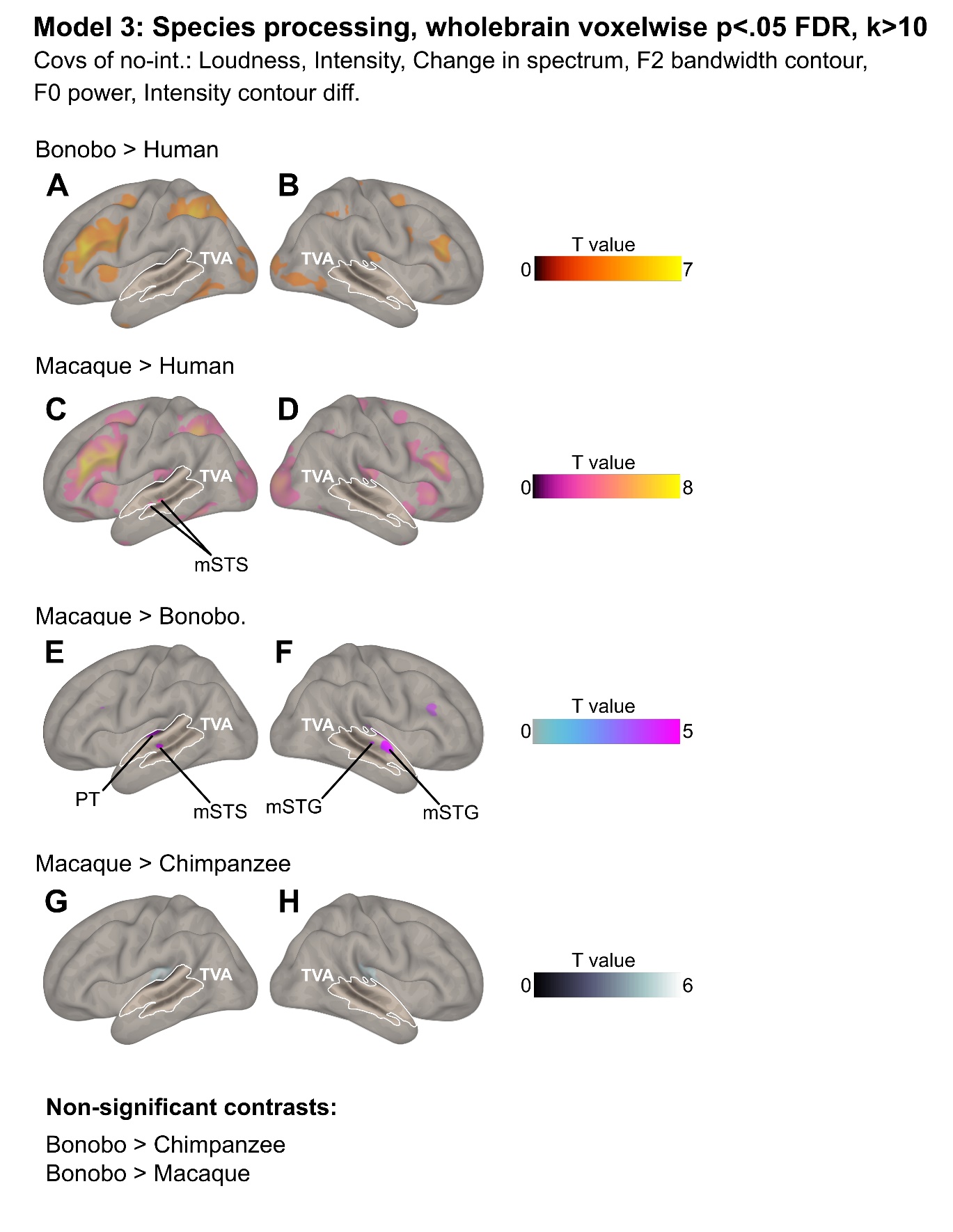

**Fig.S5: Wholebrain additional activations for model 3.** Enhanced wholebrain activity for (**A,B**) bonobo compared to human, (**C,D**) macaque compared to human, (**E,F**) macaque compared to bonobo and (**G,H**) macaque compared to chimpanzee vocalizations. Data corrected for multiple comparison using wholebrain voxelwise false discovery rate (FDR) at a threshold of *p*<.05, k=10. TVA: sample-specific (N=23) temporal voice areas; mSTG: mid superior temporal gyrus; mSTS: mid superior temporal sulcus; PT: planum temporale.

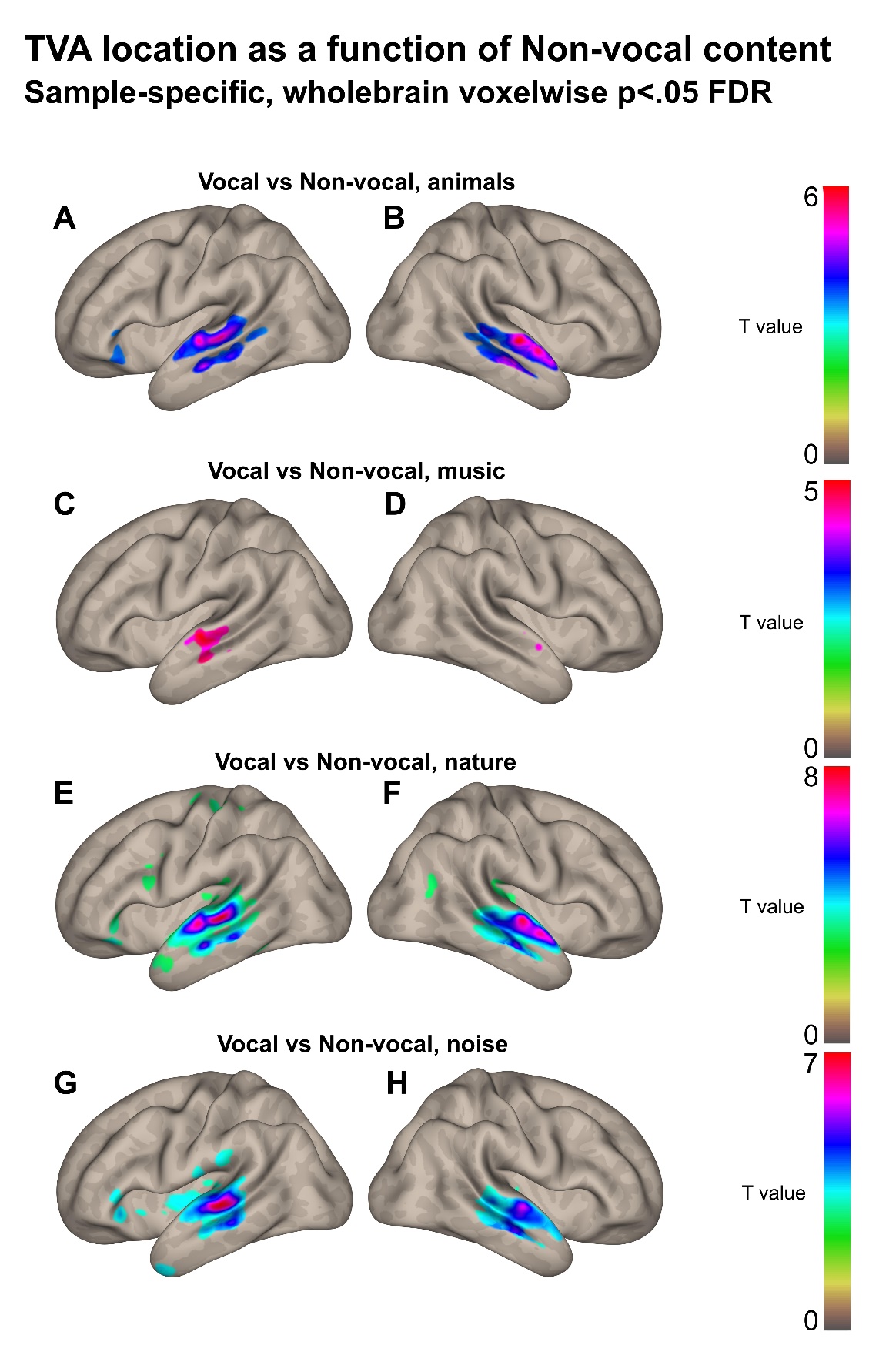

**Fig.S6: Temporal voice areas locations and subregions as a function of the type of non-vocal material, sample-specific .** Enhanced wholebrain activity for voice compared to non-voice stimuli in the main sample of the study (N=23), when non-vocal auditory material is animal sounds (**A,B**), music (**C,D**), nature sounds (**E,F**) and artificial noises (**G,H**). Data corrected for multiple comparison using wholebrain voxelwise false discovery rate (FDR) at a threshold of *p*<.05. TVA: temporal voice areas.

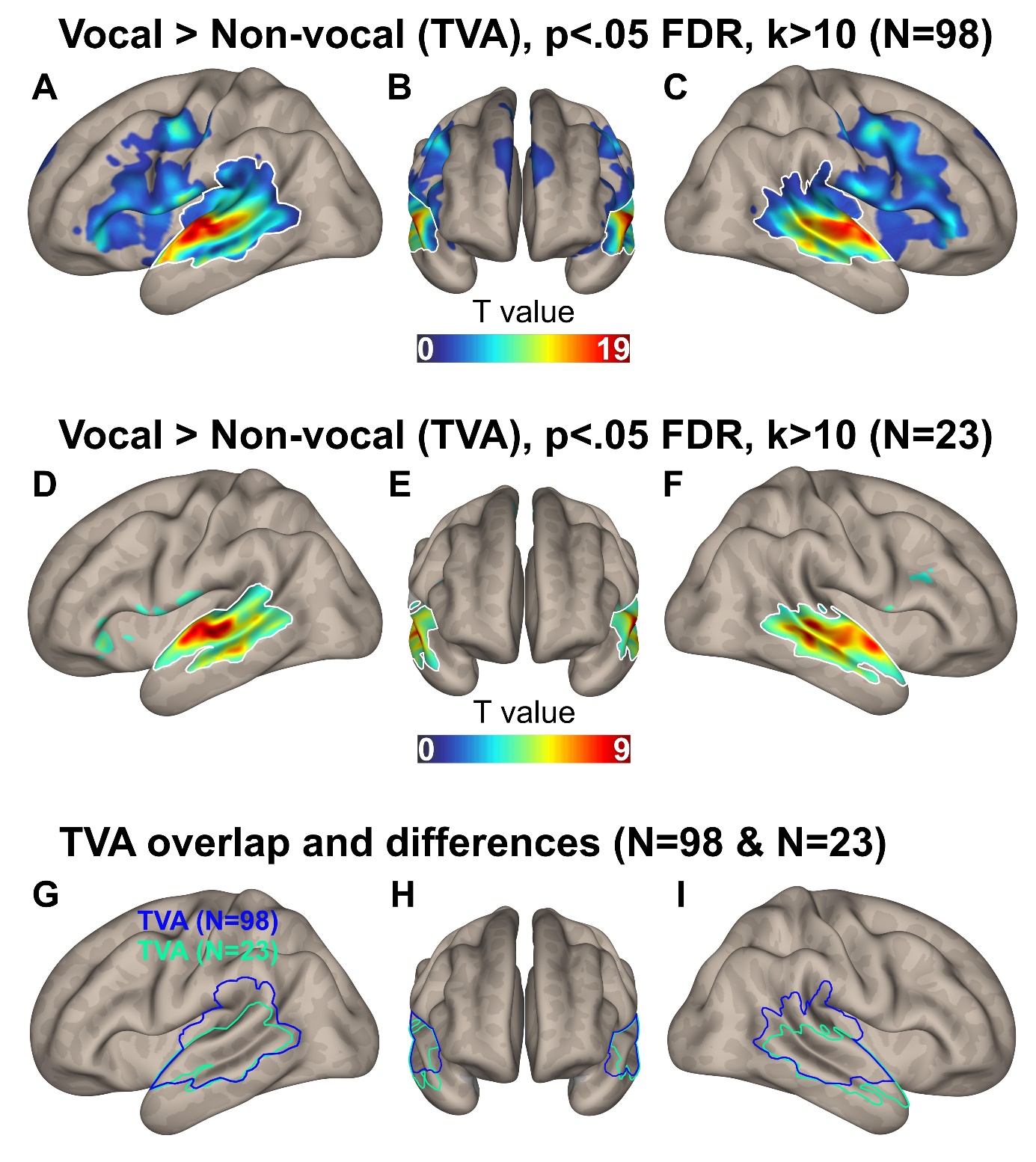

**Fig.S7: Temporal voice areas for the present study.** Enhanced wholebrain activity for voice compared to non-voice stimuli in the main sample of the study (**A,B,C,** N=23), in an independent sample of N=98 participants (**D,E,F**) and the overlap between these two samples (**G,H,I**). Data corrected for multiple comparison using wholebrain voxelwise false discovery rate (FDR) at a threshold of *p*<.05, k=10 voxels minimum per cluster. TVA: temporal voice areas.

**Table S1: Acoustic parameters (N=16) selected among a total of 88 parameters by a General Discriminant Analysis with r<.70 for the assessment of acoustic distance between species.**

| **Acoustic parameter label** | **Acoustic parameter name** |
| --- | --- |
| mfcc4_sma3_amean | Arithmetic mean of the contour of the melodic-frequency cepstral coefficient 4 |
| spectralFlux_sma3_amean***** | Arithmetic mean of the contour of the frequency with most flux around it (change in spectrum) |
| spectralFlux_sma3_stddevNorm | Normalized standard deviation of the contour of the frequency with most flux around it (change in spectrum) |
| equivalentSoundLevel_dBp***** | Conversion of power to decibel of the root mean square of the sound energy |
| slopeV0-500_sma3nz_amean | Arithmetic mean of the slope of timeframe 0-500 |
| mfcc1_sma3_amean | Arithmetic mean of the contour of the melodic-frequency cepstral coefficient 1 |
| F2amplitudeLogRelF0_sma3nz_amean | Arithmetic mean of the second formant amplitude power log relative to fundamental frequency |
| F2amplitudeLogRelF0_sma3nz_stddevNorm | Normalized standard deviation of the contour of the second formant amplitude power log relative to fundamental frequency |
| logRelF0-H1-H2_sma3nz_amean******* | Arithmetic mean of power log relative to fundamental frequency of the harmonic one and two |
| shimmerLocaldB_sma3nz_stddevNorm | Normalized standard deviation of the contour of the relative amplitude difference in local dB |
| loudness_sma3_amean***** | Arithmetic mean of the contour of the sum of the auditory spectrum |
| loudnessPeaksPerSec | Rate of peaks per second of the sum of the auditory spectrum |
| F1amplitudeLogRelF0_sma3nz_stddevNorm | Normalized standard deviation of the contour of the first formant amplitude power log relative to fundamental frequency |
| F2bandwidth_sma3nz_amean****** | Arithmetic mean of the contour of the bandwidth of the second formant |
| shimmerLocaldB_sma3nz_amean******* | Arithmetic mean of the contour of the relative amplitude difference in local dB |
| mfcc2V_sma3nz_stddevNorm | Normalized standard deviation of the contour of the melodic-frequency cepstral coefficient 2 |

**sma3:** smoothed by moving average filter with window length of 3; **sma2nz:** smoothed by moving average filter with window length of 3 for non-zero values. Saturation ([r > .70] and [r < -.70]) of factor: 1*****, 2******, 3*******.

**Table S2: Mahalanobis distances estimates, standard errors (SE) and confidence intervals (lower CI: 2.5% and upper CI: 97.5%) for the analyses using generalized linear mixed modelling of factor interaction Species * Social context for acoustic Mahalanobis distances.**

| **Term** | **Estimate** | **SE** | **Lower CI** | **Upper CI** |
| --- | --- | --- | --- | --- |
| *Agonistic ‘threat’* | | | | |
| From human to bonobo | -0.734 | 6.912 | -13.53 | 12.06 |
| From human to chimpanzee | -15.271 | 6.912 | -28.07 | -2.47 |
| From human to macaque | 20.023 | 6.912 | 7.222 | 32.822 |
| *Agonistic ‘distress’* | | | | |
| From human to bonobo | -3.037 | 6.912 | -15.836 | 9.763 |
| From human to chimpanzee | -19.491 | 6.912 | -32.291 | -6.691 |
| From human to macaque | 29.560 | 6.912 | 16.76 | 45.349 |
| *Affiliative ‘positive’* | | | | |
| From human to bonobo | 1.912 | 6.912 | -10.888 | 14.711 |
| From human to chimpanzee | -17.373 | 6.912 | -30.173 | -4.573 |
| From human to macaque | -38.699 | 6.912 | -51.499 | -25.899 |

**Table S3:** Activations, cluster size and coordinates for each contrast of interest of model 1 (mean of vocalization fundamental frequency and energy as trial-level covariates of no-interest) in the sample-specific temporal voice areas, wholebrain voxelwise *p*<.05 FDR corrected, k>10.

MNI coordinates

Region label	Hemisphere	X	Y	Z	T value	Cluster size (voxels)

**Chimpanzee > Human, Bonobo, Macaque**

Superior temporal gyrus ant^1^	L	-52	-2	-12	4.84	91

*Superior temporal gyrus mid*	L	-50	-8	-12	4.12

Superior temporal gyrus ant^2^	R	54	0	-12	3.63	18

**Chimpanzee > Bonobo, Macaque**

Superior temporal gyrus ant^3^	R	56	-2	-8	4.89	74

*Superior temporal gyrus mid*	L	-50	-8	-12	4.12

Superior temporal gyrus ant^5^	L	-52	-2	-12	5.18	78

**Chimpanzee > Human**

Superior temporal gyrus ant^4^	L	-50	-4	-12	3.36	74

**Human > Chimpanzee**

Supramarginal gyrus	R	56	-42	28	8.41	5941

*Superior temporal gyrus mid	R	56	-10	2	8.09*

*Superior temporal gyrus post	R	58	-46	14	6.76*

*Superior temporal sulcus post	R	66	-32	0	6.50*

*Middle temporal gyrus mid	R	68	-20	-10	5.52*

*Superior temporal sulcus ant	R	66	-14	-6	5.18*

*Middle temporal gyrus ant	R	60	4	-20	4.45*

Supramarginal gyrus	L	-56	-40	34	7.51	6109

*Superior temporal gyrus mid	L	-50	-18	4	7.37*

*Superior temporal gyrus post	L	-56	-52	18	6.53*

*Middle temporal gyrus mid	L	-64	-22	-10	6.10*

*Middle temporal gyrus post	L	-62	-44	-6	4.87*

*Superior temporal sulcus mid	L	-54	-32	-4	4.76*

ant: anterior; mid: central part; post: posterior.

^1-5^Figure S1 cluster labels

**Table S4:** Activations, cluster size and coordinates for each contrast of interest of model 2 (inter-species vocalization acoustic distance as trial-level covariate of no-interest) in the sample-specific temporal voice areas, wholebrain voxelwise *p*<.05 FDR corrected, k>10.

MNI coordinates

Region label	Hemisphere	X	Y	Z	T value	Cluster size (voxels)

**Chimpanzee > Human, Bonobo, Macaque**

Superior temporal gyrus ant^6^	L	-52	-2	-12	4.75	74

**Chimpanzee > Bonobo, Macaque**

Superior temporal gyrus ant^7^	L	-54	-2	-12	5.15	31

Superior temporal gyrus ant^9^	R	56	-2	-8	4.68	30

**Chimpanzee > Human**

Inferior temporal gyrus ant	L	-36	-2	-36	5.54	1310

*Hippocampus	L	-40	-24	-18	4.50*

*Superior temporal gyrus ant^8^	L	-50	-4	-12	3.30	49*

**Human > Chimpanzee**

Supramarginal gyrus	R	56	-42	28	8.91	6411

*Superior temporal gyrus mid	R	56	-8	2	8.62*

*Superior temporal gyrus post	R	58	-44	20	7.80*

*Middle temporal gyrus mid	R	68	-20	-10	5.76*

*Superior temporal sulcus ant	R	58	4	-20	5.00*

Supramarginal gyrus	L	-62	-48	24	7.66	6527

*Superior temporal gyrus mid	L	-50	-18	6	7.50*

*Middle temporal gyrus mid	L	-58	-32	-6	5.24*

*Superior temporal gyrus ant	L	-54	0	2	5.18*

ant: anterior; mid: central part; post: posterior.

^6-9^Figure S2 cluster labels

**Table S5:** Acoustic differences between chimpanzee and bonobo calls, per Context, in a dedicated analysis using eGeMaps parameters for these two species.

| **PC** | **Eigenvalue** | **Variance Explained (%)** | **Cumulative Variance (%)** | **GeMAPS Feature (raw)** | **Short Name** | **Dominant Loading** |
| --- | --- | --- | --- | --- | --- | --- |
| PC1 | 2,992 | 12.47 | 12.47 | F0semitoneFrom27.5Hz_sma3nz_pctlrange0.2 | F0 Range (semitones) | +0.349 |
| PC2 | 2,281 | 9.50 | 21.97 | slopeV500.1500_sma3nz_stddevNorm | Spectral Slope 500–1500 Hz Variability | +0.441 |
| PC3 | 2,129 | 8.87 | 30.84 | F1frequency_sma3nz_stddevNorm | F1 Frequency Variability | +0.357 |
| PC4 | 1,771 | 7.38 | 38.22 | StddevUnvoicedSegmentLength | Unvoiced Segment Duration SD | +0.432 |
| PC5 | 1,639 | 6.83 | 45.05 | HNRdBACF_sma3nz_stddevNorm | HNR Variability | -0.339 |
| PC6 | 1,459 | 6.08 | 51.13 | slopeV0.500_sma3nz_amean | Spectral Slope 0–500 Hz (mean) | +0.474 |
| PC7 | 1,394 | 5.81 | 56.94 | F1bandwidth_sma3nz_stddevNorm | F1 Bandwidth Variability | +0.414 |
| PC8 | 1,253 | 5.22 | 62.16 | F0semitoneFrom27.5Hz_sma3nz_stddevRisingSlope | F0 Rising Slope Variability | -0.403 |
| PC9 | 1,22 | 5.09 | 67.24 | F2amplitudeLogRelF0_sma3nz_stddevNorm | F2 Amplitude Variability (rel. F0) | +0.387 |
| PC10 | 1,051 | 4.38 | 71.62 | slopeV0.500_sma3nz_stddevNorm | Spectral Slope 0–500 Hz Variability | +0.358 |
| PC11 | 0,984 | 4.10 | 75.72 | F3amplitudeLogRelF0_sma3nz_stddevNorm | F3 Amplitude Variability (rel. F0) | +0.520 |
| PC12 | 0,906 | 3.77 | 79.50 | jitterLocal_sma3nz_stddevNorm | Jitter Variability | +0.407 |
| PC13 | 0,88 | 3.67 | 83.16 | logRelF0.H1.A3_sma3nz_stddevNorm | H1–A3 Amplitude Ratio Variability | +0.470 |

PC : principal component ; F0 : fundamental frequency ; F1 : first formant ; F2 : second format ; F3 : third formant ; stddev : standard deviation ; Norm : normalized.
